# *SpacerPlacer*: Ancestral reconstruction of CRISPR arrays reveals the evolutionary dynamics of spacer deletions

**DOI:** 10.1101/2024.02.20.581079

**Authors:** Axel Fehrenbach, Alexander Mitrofanov, Omer S. Alkhnbashi, Rolf Backofen, Franz Baumdicker

**Affiliations:** Cluster of Excellence “Controlling Microbes to Fight Infections”, Mathematical and Computational Population Genetics, University of Tübingen, 72076 Tübingen, Germany; Institute for Bioinformatics and Medical Informatics (IBMI), University of Tübingen, 72076 Tübingen, Germany; Bioinformatics group, Department of Computer Science, University of Freiburg, 79085 Freiburg, Germany; Center for Applied and Translational Genomics (CATG), Mohammed Bin Rashid University of Medicine and Health Sciences (MBRU), Dubai Healthcare City, 505055 Dubai, United Arab Emirates; College of Medicine, Mohammed Bin Rashid University of Medicine and Health Sciences (MBRU), Dubai Healthcare City, 505055 Dubai, United Arab Emirates; Signalling Research Centres BIOSS and CIBSS, University of Freiburg, 79085 Freiburg, Germany

**Keywords:** CRISPR-Cas, bacterial genomics, spacer array, ancestral reconstruction, spacer gain and loss, likelihood functions, microbiomes, CRISPR typing

## Abstract

Bacteria employ CRISPR-Cas systems for defense by integrating invader-derived sequences, termed spacers, into the CRISPR array, which constitutes an immunity memory. While spacer deletions occur randomly across the array, newly acquired spacers are predominantly integrated at the leader end. Consequently, spacer arrays can be used to derive the chronology of spacer insertions.

Reconstruction of ancestral spacer acquisitions and deletions could help unravel the coevolution of phages and bacteria, the evolutionary dynamics in microbiomes, or track pathogens. However, standard reconstruction methods produce misleading results by overlooking insertion order and joint deletions of spacers.

Here, we present SpacerPlacer, a maximum likelihood-based ancestral reconstruction approach for CRISPR array evolution. We used SpacerPlacer to reconstruct and investigate ancestral deletion events of 4565 CRISPR arrays, revealing that spacer deletions occur 374 times more frequently than mutations and are regularly deleted jointly, with an average of 2.7 spacers. Surprisingly, we observed a decrease in the spacer deletion frequency towards *both* ends of the reconstructed arrays. While the resulting trailer-end conservation is commonly observed, a reduced deletion frequency is now also detectable towards the variable leader end. Finally, our results point to the hypothesis that frequent loss of recently acquired spacers may provide a selective advantage.

## Introduction

Bacteria and Archaea are frequently exposed to phage and plasmid invasions. As a countermeasure, an enormous number of immune mechanisms have evolved in prokaryotes [1, 2]. A very prominent system is the adaptive defense system CRISPR-Cas, Clustered Regularly Interspaced Repeats (CRISPR) with CRISPR-associated genes (Cas), which provides immunity against phages, plasmids, and other mobile genetic elements [3, 4]. A variety of CRISPR-Cas systems have been found and categorized into two classes with a multitude of types and subtypes [5, 6] depending on the composition and structure of the found Cas-cassettes. The CRISPR-Cas system defends against an invader by acquiring a so called spacer which corresponds to a small snippet, called a protospacer, of the invading genetic element. These spacers are inserted into the CRISPR spacer array [4, 7]. During subsequent invasions, the individual spacers are then expressed as guide RNAs and combined with Cas proteins. These Cas-complexes are then able to find the corresponding protospacers and interfere with or degrade the target depending on the involved Cas-complex [6, 8]. While the mechanisms of interference are highly subtype specific and depend on the specific set of Cas proteins involved, many of the Cas proteins responsible for adaptation (e.g. Cas1 and Cas2) are widespread and more conserved. Nevertheless, subtype-specific differences in adaptation exist.

### CRISPR spacer array evolution

The adaptation process, i.e. the acquisition of spacers, is an essential part of most CRISPR systems. Spacer sequences are inserted into an array separated by repeats [4, 7]. Remarkably, insertion of new spacers is almost always facilitated at one end of a CRISPR array [7, 9]. Thus, the order of spacers in a CRISPR array provides a timeline of the phages and plasmids encountered, identified by the corresponding protospacer sequences.

As part of the genome, acquired CRISPR spacers are inherited across generations, but can also be deleted [10]. The deletion of DNA has been shown to be weakly beneficial since replication of longer genomes carries a cost for the cell. In case of CRISPR arrays, deletion of spacers can not only result in immunity loss but carries an additional importance since the effectiveness of the remaining spacers depends on their position and the length of the array [11]. While in adaptation, e.g. factors like Cas1, Cas2 and the leader sequence are well studied, the source for deletions is not yet fully understood. Hypotheses include that spacer deletion occurs by misalignment between repeats during replication [11, 12], or with the addition of a new spacer [13]. Spacer acquisition and deletion are frequent and thus constantly modify the CRISPR arrays within the population, resulting in observable short-term dynamics formed by the co-evolution with foreign genetic elements. The ordered timeline of spacer acquisitions not only constrains the population histories but also links to the targeted co-evolving phages and plasmids. Furthermore, by studying the acquisition and deletion processes and comparing them to neutral evolutionary models we are able to investigate, if selection effects occur. Finally, the arrays are relatively short in length and can be reliably identified due to their repeat structure [14–17].

In principle, CRISPR arrays could thus be particularly useful for investigating evolutionary relationships at high resolution, tracking and typing samples in epidemiological outbreaks, understanding the dynamic ecology of microbiomes, and analyzing biotechnological and pharmaceutical biocultures [17–24]. Recently, López-Beltrán et al. observed that the abundance of phages varies depending on the position of their corresponding spacers in the array [25]. Thus, we are particularly interested, if differences in selection strengths depending on the age of spacers, that is their position in the array, can be inferred from deletion events. However, a reconstruction tool, as presented here, is needed to correctly determine ancestral spacer insertion and deletion events.

### CRISPR spacer array evolution challenges traditional phylogenetic reconstruction methods

Reconstructing bacterial evolution from CRISPR arrays faces the problem that the number of spacers in an array is small, and that polarized acquisitions and blockwise deletions generate stochastic dependencies between different evolutionary spacer events. This makes it difficult to use standard phylogenetic methods for DNA sequences, which neglect the specifics of CRISPR evolution, resulting in avoidable errors.

While a variety of visualization tools for CRISPR spacer arrays have been developed [8, 26–28], reconstruction methods that take into account the specifics of CRISPR array evolution are still rare. An exception is the CRISPR Comparison Toolkit (CCTK) introduced by Collins and Whitaker [8], which contains a maximum parsimony approach to reconstruct ancestral spacer arrays by first aligning the arrays using the Needleman-Wunsch algorithm, and then inferring ancestral arrays using empirically determined parsimony costs. However, to go beyond ancestral reconstruction, it is crucial to employ explicit evolutionary models. These models incorporate interpretable parameters, allowing for a more comprehensive analysis of the corresponding molecular mechanisms and evolutionary processes.

### SpacerPlacer allows to investigate spacer array evolution

Here, we introduce *SpacerPlacer* a new parametric likelihood-based approach tailored for the reconstruction of ancestral spacer insertions and deletions. *SpacerPlacer* reconstructs ancestral states while considering the unique features of CRISPR arrays and their polarized insertion order through an evolutionary model. The advantage of using a probabilistic approach is that we can compare different evolutionary models for CRISPR spacer arrays and investigate the corresponding molecular mechanisms of spacer acquisitions and deletions, as well as selection pressures, in more detail. One major benefit of this model-based approach is that it - in addition to the reconstruction - allows to estimate the underlying evolutionary parameters. This enables us to infer the distribution of the number of jointly lost adjacent spacers, detect potential differences between CRISPR-Cas types, and test model assumptions using likelihood ratio tests.

We applied *SpacerPlacer* on all suitable high-confidence CRISPR arrays from the CRISPRCasdb database [29]. Despite the variability of subtypes, our analysis did not find significant differences in deletion rates and lengths between the subtypes considered here, which points to a Cas-independent process for deletion. We find that the median deletion rate per spacer is 374 times higher than the mutation rate per site. Since spacers and their repeats are fairly short [12, 30], we conclude that the evolution of CRISPR arrays by acquisitions and deletions is much faster than the mutation of the spacer sequences themselves.

We then investigate the reconstructed spacer deletions in more detail. To this end, we performed a likelihood ratio test and found that spacers are indeed regularly jointly deleted. Using our probabilistic framework, we estimate the mean length of these joint deletions as 2.7 adjacent spacers.

We also investigated how the distribution of the deletion frequency changes with respect to the position within the array. We find that deletion frequencies are decreased towards both boundaries of the array. This boundary effect is consistent with the conjecture that spacer deletions are mainly caused by repeat misalignment.

Moreover, by comparing empirical and simulated deletion frequencies, we find indications that either selection increases the deletion frequency of the first spacer, or mutations in the last repeat reduce the deletion frequency of the last spacer, or both.

## Methods

### Spacer array data preparation

To run *SpacerPlacer* we require a group *G* of completely sequenced related CRISPR arrays with consecutively ordered spacers and a respective phylogenetic tree *𝒯*.

Ideally, this tree should be provided by the user, but *SpacerPlacer* provides a heuristic method to infer a suitable tree *𝒯* from the spacer arrays itself.

Each unique spacer DNA sequence in *G* is labeled with an individual natural number. We omit an illustration here, but simple examples of a group *G* can be found in Figure 2 and Figure 3a). Obviously, the arrays in *G* should share some spacers but not all be identical. So, we restricted the analysis to such groups.

**Fig. 1.**
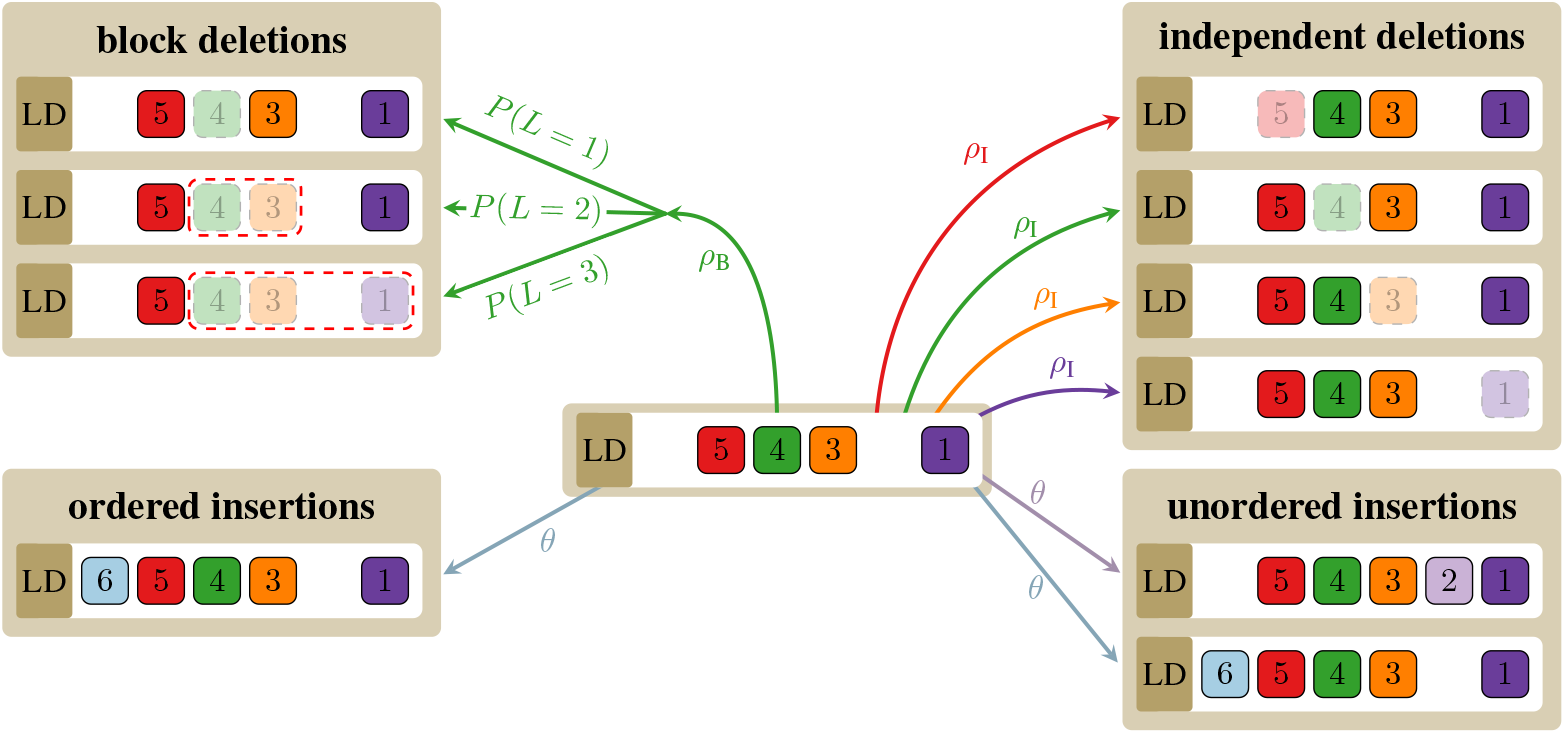
Different deletion and insertion models. The figure shows how aligned spacer arrays evolve in different models used within *SpacerPlacer*. Each colored and numbered box represents a different spacer. Unordered insertions can occur at any unoccupied position within the array (bottom right). Within the ordered insertion model (bottom left), spacers can only be acquired directly after the leader (indicated by LD) in front of the first existing spacer. In the independent deletion model (top right), each spacer can be deleted individually with rate *ρ*_*I*_. Block deletions (top left) occur at each (occupied) position at rate *ρ*_*B*_ and *L ≥* 1 spacers are deleted jointly (including the spacer hit by the event), where *L* is geometrically distributed with mean *α*. For simplicity, we show only one block deletion event. The color of the arrows indicates at which spacer position an event was initiated. The most realistic model includes ordered insertions and block deletions (left side), while models with independent deletions and unordered insertions (right side) serve as a reference and speed up parts of the reconstruction pipeline.

**Fig. 2.**
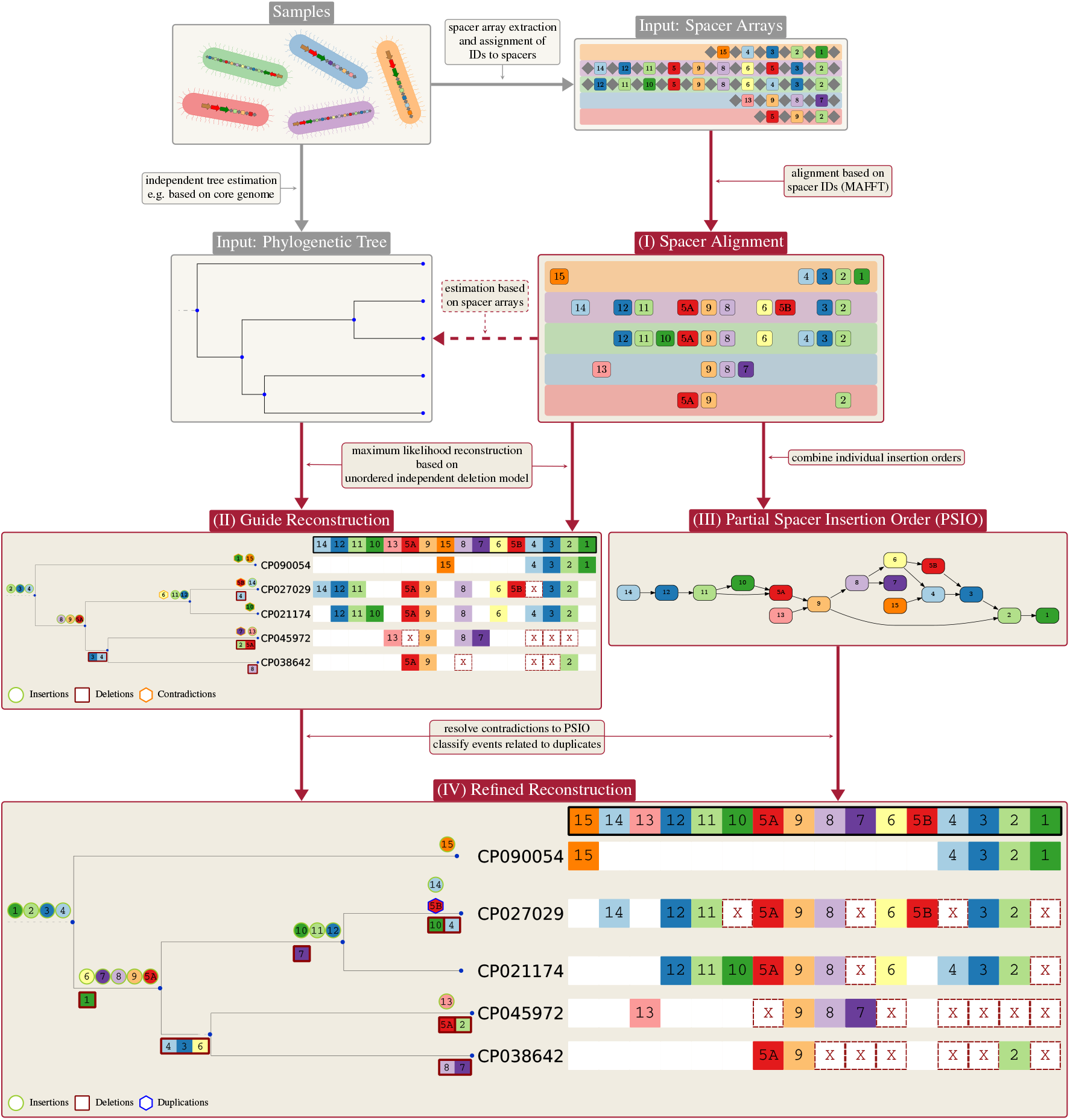
Workflow of *SpacerPlacer* : ancestral reconstruction. To prepare for the *SpacerPlacer* analysis the first step is to extract the CRISPR arrays from their sample genomes and reconstruct a phylogenetic tree, e.g. based on genome alignments of the samples. Then each unique spacer (cluster) gets assigned an ID. This step should generally be done by the user, but *SpacerPlacer* includes ID assignment for spacer arrays in a CRISPRCasdb/Finder format. As a first step, *SpacerPlacer* aligns the spacer arrays in their ID form. The spacer alignment optionally allows us to reconstruct an alternative phylogenetic tree. Then *SpacerPlacer* builds the partial spacer insertion order (PSIO) and performs a guide reconstruction along the provided tree. This is followed by refining the reconstruction using the PSIO, resulting in a finished reconstruction of ancestral acquisition, deletion and more complex events of the considered group. Preparation steps are shown in gray boxes and steps done by *SpacerPlacer* are shown in red boxes. The IDs can be chosen arbitrarily. Here, we use an ID assignment that makes the alignment more intuitively readable. The group portrayed here are CRISPR arrays of Cas type I-B from *Listeria monocytogenes* extracted from CRISPRCasdb. The tree shown is estimated based on the CRISPR arrays with *SpacerPlacer*.

**Fig. 3.**
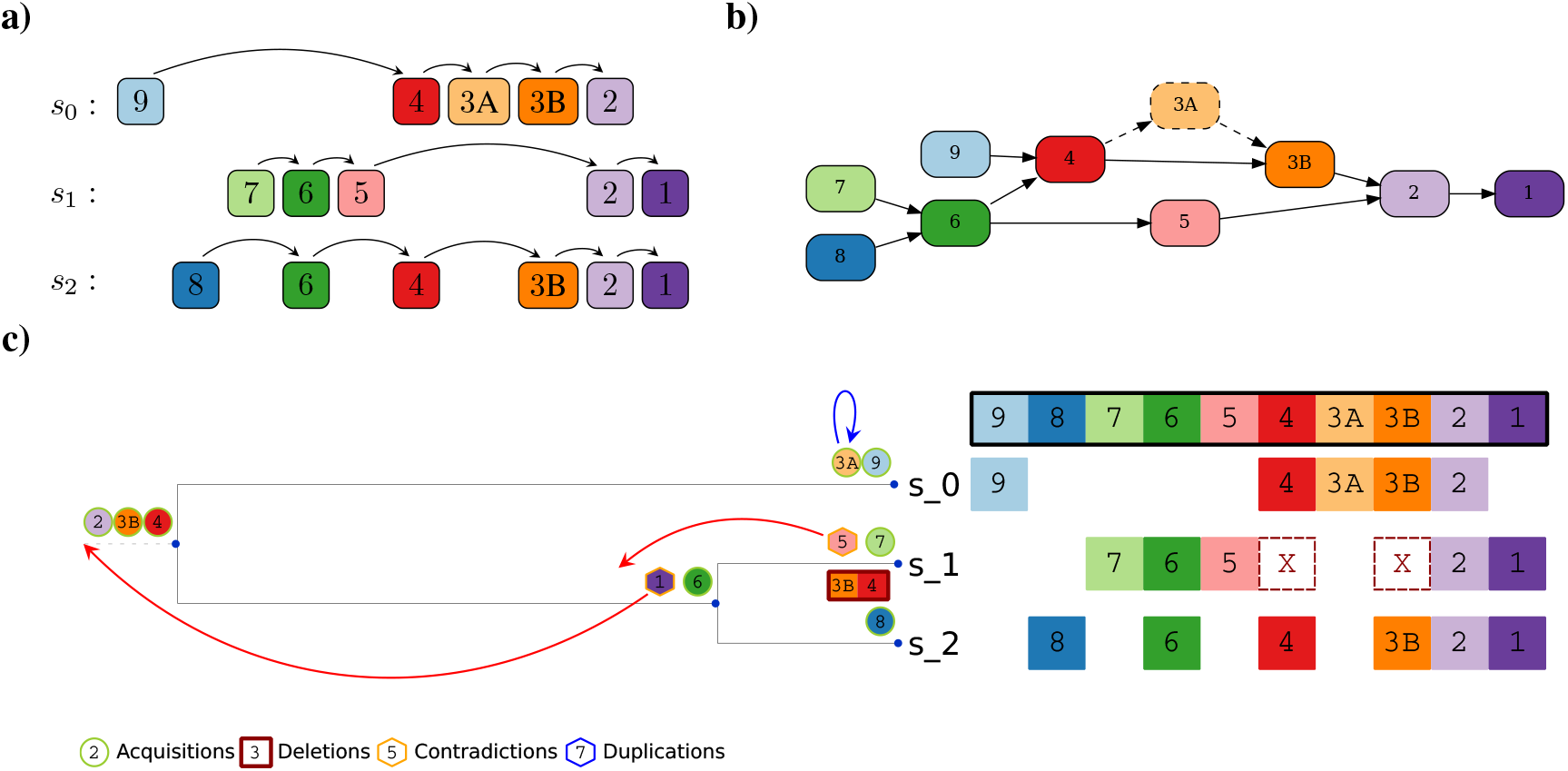
Partial Spacer Insertion Order based refinement of the guide reconstruction. **a)** Relabeled multiple spacer alignment of a group with 3 arrays *s*_0_, *s*_1_ and *s*_2_, together with their induced orders. Only one step relationships are indicated. Spacers 3A and 3B are relabeled duplicates of each other. **b)** Partial Spacer Insertion Order (PSIO) as induced by the spacer array group in a). The spacer 3A is categorized as a potential duplication event occurring between 3A and 3B. Thus, it is removed from the PSIO for the refined reconstruction (dashed), and contradictions due to 3A are ignored. **c)** Illustration of the guide reconstruction and the PSIO-based refinement process. The acquisition event of 5 contradicts the acquisition of 6 in the parent branch. Similarly, the insertion of 1 is a contradiction to 2 (or 3, 4). Both contradictions are solved in the refinement process by fixing the spacer to be present at the parent node, as indicated by the red arrows. Spacer 3A is classified as a duplication event, since 3A is located in the middle of the array and spacer 3B is already present on this branch, indicated by the blue arrow.

#### Database

CRISPRCasdb is a freely accessible database that contains the CRISPR array predictions of “CRISPRCas-Finder” [31] on whole genome assemblies collected from GenBank. For the construction of the spacer database, we extracted spacer sequences from CRISPRCasdb [29] in July 2022. We only consider predictions of the highest evidence level (4), resulting in a total of 7352 repeats and 32624 arrays.

#### Clustering spacers based on sequence homology

After extraction, we clustered the spacers into sets that we treat as identical, and have likely undergone only mutations. Each spacer in the CRISPR arrays was relabeled according to its respective cluster label. The relabeled CRISPR arrays serve as input for *SpacerPlacer*.

The clustering was performed individually for each identified consensus repeat. The procedure encompassed several stages: Initially, all CRISPR arrays containing the specific consensus repeat were selected. Subsequently, spacers associated with these arrays were aggregated. Hierarchical clustering was then applied to these spacers using the following algorithm: Initially, each spacer (combined with its reverse complement) was treated as an independent cluster. In subsequent iterations, the clusters were progressively merged if they contained spacer pairs that differed by a single Levenshtein editing operation (substitution, insertion, deletion).

In theory this liberal clustering method could result in huge clusters. However, in practice, we find clusters to be small. For all 53965 repeats on CRISPRCasdb, more than 97% had solely identical spacer sequences in the associated clusters. For the remaining 1426 repeats, the largest spacer cluster has size 11, but most of them had size 2.

#### Identification of sets of closely related CRISPR arrays

We further subdivided the pool of closely related arrays with the highest confidence level according to their repeat sequence. For all repeats, we checked if there was an overlap between reverse complement repeats and reverse complemented spacer arrays.

CRISPR arrays were assigned their Cas type according to the closest Cas-cassette found within the genome. Arrays with no close (< 10000 bp distance) Cas-cassette were removed unless only one Cas-cassette was found within the genome. Groups were further categorized by genus and Cas type. Finally, the groups were further subdivided by pairwise overlap of the spacers. Each spacer array has to have at least one spacer overlap with any of the other arrays in a group to be included. We excluded groups that had less than three arrays or lacked spacer diversity between arrays. Thus, within each group, each sample has the same repeat, genus, and Cas type. For further details and a breakdown of the number of groups and arrays within each step see Supplementary Note 1A.

The final dataset with core gene trees fed into *SpacerPlacer* contains 518 groups with 5934 arrays in total. Of these, 333 groups (4565 arrays) contain at least one reconstructed deletion. We refer to these groups as the “CRISPRCasdb dataset”. A breakdown of the Cas types and genera found in the CRISPRCasdb dataset is shown in Supplementary Figure S6 and S7, respectively.

#### Multiple spacer array alignment

The order of spacers in different arrays, and sometimes even within a single spacer array, can be inconsistent and produce contradictions if we assume polarized insertions. For instance, if a spacer occurs twice in array A and only once in array B, it is unclear which of the duplicated spacers in array A corresponds to the spacer in array B. Another type of inconsistency can arise if spacer 1 follows spacer 2 in array A but spacer 2 follows spacer 1 in array B. This is clearly not realizable through a single acquisition.

To identify and group such candidate spacers, *SpacerPlacer* performs a Multiple Spacer Array Alignment (MSAA) on each group of spacer arrays. We then identify the most plausible matching spacers according to their positions in the MSAA. We align spacer arrays solely based on the spacer labels. Note that we do *not* align the spacer DNA sequences. For the multiple spacer alignment we use the tool MAFFT [32], since MAFFT can also align arbitrary characters (the spacer labels in our case) instead of nucleotide and protein sequences. *SpacerPlacer* tweaks the MAFFT parameters and uses low gap penalties and a very high mismatch penalty, to prevent spacer mismatches, favoring gaps instead. More details about this process can be found in Supplementary Note 2.

#### Tree estimation

The quality of the underlying phylogenetic tree seriously affects both the ancestral spacer reconstruction and the parameter estimates. In general, we suggest using whole genome trees, if available, to reduce the dependency between tree estimation and array reconstruction, especially, if rate estimation is desired.

For each group of spacer arrays, we reconstructed a phylogenetic tree, independent of the arrays, based on the single-copy core genes of the corresponding sample. We used Bakta to annotate genomes [33], the pangenome analysis tool panX [34], and IQ-TREE [35] to reconstruct trees. Throughout this work, we used these core genome trees unless otherwise stated.

We did not find evidence indicating that our median estimates were affected by potential biases introduced by using core genome trees for CRISPR arrays where horizontal transfer might have occurred. Furthermore, we removed samples with multiple arrays from our analysis to avoid the inability of core genome-based trees to differentiate between them.

As an alternative for cases where no plausible independent tree is available, *SpacerPlacer* offers the option to use basic unweighted pair group method with arithmetic mean (UPGMA) or neighbor joining (NJ) approaches to infer trees directly from CRISPR arrays. For this, we developed a distance function for spacer arrays that takes into account the specifics of CRISPR array evolution, including blockwise deletions and acquisition order. The details of the distance function can be found in Supplementary Note 3.

#### (Re-)orientation of spacer arrays

The orientation of spacer arrays was chosen according to a comparison between the likelihood of the “forward” reconstruction of the group, i.e. the orientation supplied by CRISPRCasdb, and the likelihood of the “reverse” reconstruction, where the MSAA of the group was reversed. If no clear distinction, i.e. a large margin between likelihoods was found, the orientation provided by CRISPRCasdb was used.

### Simulations

The simulation of spacer acquisition and deletion events along a simulated tree was done similarly to previous approaches [36, 37].

We start at the root of the corresponding phylogeny, with spacer arrays with a random length that is distributed according to the stationary length distribution of the respective evolutionary model. The array is then evolved along the provided tree by sampling insertions and deletion events according to an independent or block deletion model with polarized insertions. The underlying trees were simulated according to the coalescent, if not indicated otherwise. More details can be found in Supplementary Note 1B.

### Estimation of block deletion length in experimental data

Naturally, the actual process that determines the number of jointly lost spacers is more complex than a geometrically distributed random variable, but this simplification provides an efficient and precise computation of the likelihood function in *SpacerPlacer*. However, we found statistical evidence that the lengths of experimentally derived spacer deletion blocks are well approximated by a geometric distribution.

To test the approximation of geometrically distributed lengths of block deletions we performed a goodness of fit test (Kolmogorov-Smirnov) on the empirical distribution inferred by Lopez-Sanchez et al. [38]. They selected against an individual spacer (number 258) and experimentally determined the neighboring spacers that were deleted jointly with this spacer (Figure 3 in [38]). Since the time span of the experiment was very short and only a few (2) concurrent spacer deletions occurred, we assume the deletions of and around spacer 258 to be the result of single events. Therefore, we conditioned on a guaranteed deletion of spacer 258 to estimate the expected length of deletions.

## Results

### A new probabilistic model for CRISPR array evolution

#### Underlying probabilistic model

We will introduce different models, all Markov processes along a given (estimated) phylogenetic tree *𝒯* (note that we adapt the naming conventions of previous studies [36, 37]).

The state space in our models corresponds to the alignment of all (observed) spacers. At each branch, the spacer array is described by a presence-absence pattern, where each spacer corresponds to a position in the input alignment.

Spacer insertion and deletion events occur along the branches of *𝒯* according to a Poisson point process. Thus, each of the models is characterized by rate-of-occurrence parameters.

The main differences between the models are the possible positions along the spacer array where new spacers can be inserted and the specific characteristics of deletion events, as described in the following.

#### Ordered or unordered insertions

Despite variations in the adaptation proteins Cas1 and Cas2 [5, 6, 12, 39], the polarized insertion remains nearly universal across Cas types, and we leverage this fact to improve the reconstruction in *SpacerPlacer*. Thus, we mainly consider the *ordered insertion model* corresponding to the predominant mode of insertion [7], where a spacer can be inserted only at the leader end. While the ordered insertion model is clearly the more realistic model, we also introduce a simpler *unordered insertion model* to speed up and simplify parts of the guide reconstruction. In this model, new spacers can also be inserted between existing spacers. The different events of both models are shown for an example array in Figure 1. For all models, we assume that all acquired spacers are unique. To guarantee this in the reconstruction algorithm, *SpacerPlacer* renames multiple occurrences of the same spacer and thus guarantees a unique naming of spacers.

#### Independent or blockwise deletions

In contrast to insertions, so far no active molecular mechanism modulating spacer deletions has been identified. Replication misalignment or strand/replication slippage have been proposed as the main underlying molecular mechanisms that cause spacer deletions between repeats [12]. Misalignment is believed to occur when two repeats erroneously align within a single replication fork [40]. Consequently, this process is constrained to repeats that are relatively close in proximity, which aligns well with CRISPR arrays, where spacers between repeats tend to be rather short. Furthermore, this would also allow for deletions of several adjacent spacers, which we term block deletions.

Thus, to investigate how many spacers are typically affected by a single deletion event, we contrast two different deletion models. In the *independent deletion model (IDM)* each individual spacer *s*_*i*_ (of some array *s* = (*s*_1_, *s*_2_, …, *s*_*n*_) of length *n*) can only be independently deleted with rate *ρ*_I_. In the *block deletion model (BDM)*, deletion events also arise independently at any spacer *s*_*i*_ at rate *ρ*_B_. However, a consecutive block of *L* spacers *s*_*i*_ … *s*_*i*+*L−*1_ is jointly deleted, where *L* is geometrically distributed with mean *α*. If *L* exceeds the boundary of the spacer array, the event is ignored. Note that, due to its monotonically decreasing probability mass function, longer deletions become increasingly unlikely and singular deletions remain the most likely. Further, note that setting *α* = 1 yields the independent deletion model, which allows us to use the framework of likelihood ratio tests to distinguish between the IDM and the BDM. Both models are illustrated in Figure 1.

An alternative result of replication misalignment is spacer duplication or rearrangement [41, 42], which has also been observed for spacer-repeat units in CRISPR arrays [12, 40]. Here, we do not directly incorporate such more complex events into the evolutionary models. Instead, we detect and resolve them in a separate step of the reconstruction pipeline.

### Ancestral CRISPR Spacer Array Reconstruction

#### Overview of the pipeline

*SpacerPlacer* is a parametric reconstruction and visualization tool for CRISPR spacer array evolution. An overview of the reconstruction pipeline is shown in Figure 2. Based on a set of overlapping spacer arrays and a provided phylogeny, *SpacerPlacer* proceeds in multiple steps to reconstruct ancestral spacer acquisitions, deletions, and more complex events, such as rearrangements.

I. First, spacers are represented by the labels that group almost identical spacers and a spacer array alignment is constructed based on these labels instead of aligning the DNA sequences. The spacer array alignment is described in more detail in Methods. Aligning the spacer labels prevents misalignment of different spacers. The alignment of the spacer array reveals duplicated spacers as well as candidates for more complex events.
II. The dependence between different spacer sites introduced by polarized insertion and block deletions makes computing the joint likelihoods for the reconstructions infeasible. Therefore, we first calculate preliminary ancestral states at the internal nodes of the provided phylogeny based on the less complex independent deletion model with unordered insertions.
III. The guide reconstruction does not account for the relative acquisition-time information caused by polarized insertion, which can be seen partially in the alignment of the spacer arrays. We extract this information from the MSAA into a partial spacer insertion order (PSIO) which is explained in more detail below.
IV. Finally, the guide reconstruction is refined so that the reconstructed insertions obey the partial order. This indirect inclusion of the insertion polarization is more efficient than calculations in a complex parametric model. Finally, the reconstructed events of putative duplicated spacers are categorized as duplications, rearrangements, reinsertions, or similar events. On the basis of such reconstructed insertion and deletion events, *SpacerPlacer* will also estimate the parameters of an IDM and the more complex BDM. Subsequently *SpacerPlacer* identifies the most likely model in a likelihood ratio test (Figure 4) for the dataset considered. *SpacerPlacer* is available at https://github.com/fbaumdicker/SpacerPlacer. For additional information about the implementation, used software and packages see Supplementary Note 2. The analysis of a set of spacer arrays on a standard laptop takes usually just a few seconds. The analysis of all 518 groups from CRISPRCasdb took 515.13 seconds.

**Fig. 4.**
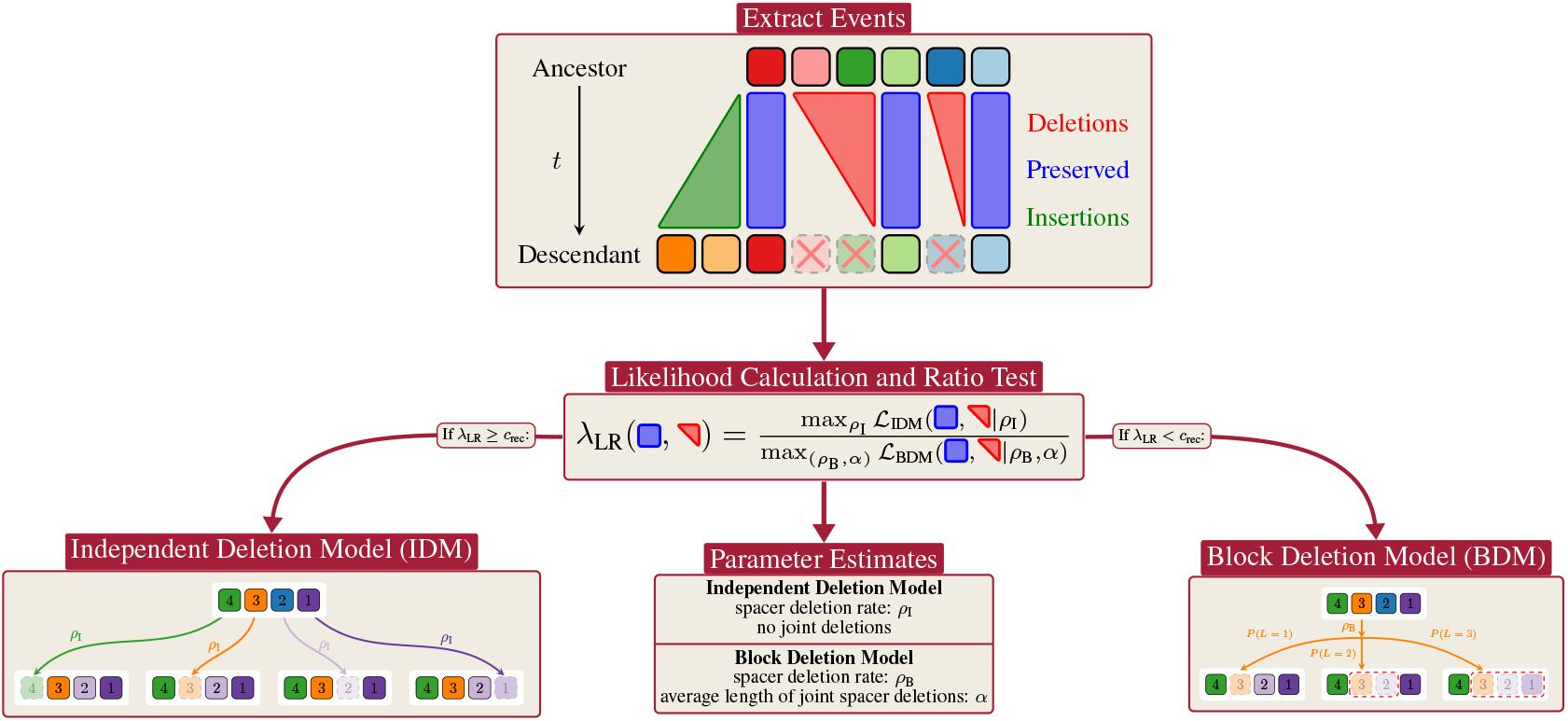
LRT workflow in SpacerPlacer. Using a parametric model allows *SpacerPlacer* to combine ancestral reconstruction with rate estimation and likelihood ratio tests. After reconstructing the ancestral states with *SpacerPlacer* we extract the deletion events and preserved spacers, as well as the lengths of the tree branches. These events are then used to generate maximum likelihood estimates for the parameters of both deletion models the IDM and the BDM. *SpacerPlacer* also uses a likelihood ratio test to identify whether the IDM can be rejected in favor of the BDM.

#### Guide reconstruction

As a first step of the ancestral spacer array reconstruction we calculate a preliminary *guide* reconstruction. As the unordered independent deletion model on spacer presence-absence profiles is a *general time reversible (GTR)* model we can use an effective dynamic programming algorithm [43] for our ancestral guide reconstruction. As multiple independent acquisitions of the same spacer are rare, we use parameters that encourage the model to prioritize multiple deletions over multiple insertions in the guide reconstruction.

The algorithm scales linearly with the number of arrays and linearly with the number of sites, in our case, the number of spacers in the MSAA [43]. Our implementation is based on *TreeTime* [44] and outlined in more detail in Supplementary Note 2.

The guide reconstruction cannot take advantage of all available information as it ignores the spacer insertion order. Consequently, it can still contradict polarized acquisition, as illustrated in Figure 3. In particular, the guide reconstruction will still underestimate the number of deletion events. However, it is a suitable starting point for the subsequent refinement steps that address these issues.

#### Partial Spacer Insertion Order graph construction

The partial order within the spacer arrays encodes the information that is generated from polarized insertions [7], one of the most consistent properties of CRISPR arrays. This has been empirically confirmed for many Cas types, except for type II-A systems, where ectopic insertions have also been found [45–47].

Each spacer array constitutes a timeline of acquired spacers, i.e., an insertion order for the acquisitions of this array. Thus, by combining the order information of all arrays in a group, we can extract additional timeline information. For example, when given two arrays, A (1-2-4-6) and B (2-3-5-6). Here, array A implies that spacer 1 was acquired after spacer 2 and array B implies spacer 2 after spacer 3. Together, we can conclude that spacer 1 was acquired after spacer 3. Note that when considering multiple spacer arrays, not all spacers need to have an order relation with each other. In the previous example, the arrays do not determine whether spacer 4 was inserted before or after spacer 5. Thus, combining the order information of multiple arrays leads only to a *partial* spacer insertion order.

The PSIO can be visualized as a graph (Figure 3b)). In this graph, each arrow indicates a “younger than” or equivalently, “was acquired after” relation. We construct the PSIO by thinning out redundant edges from the graph based on all single-array insertion orders from the MSAA. The resulting graph is a directed graph without loops as the previous MSAA step already resolved potential conflicts. *SpacerPlacer* automatically generates a visualization of the PSIO graph with graphviz [48].

#### Iterative PSIO-based refinement

ssThe guide reconstruction does not need to be consistent with the PSIO and can contradict polarized acquisition. Therefore, we iteratively refine the guide reconstruction using three straightforward steps:

1. Identify any contradictions between the insertions estimated by the ancestral reconstruction and the PSIO. If no contradictions are found, the reconstruction is complete. Otherwise, continue with step 2.
2. For each reconstructed spacer acquisition event that is in conflict with the PSIO, determine the parent node in the tree. Regardless of the most likely state calculated at this node in the guide reconstruction, fix the spacer to be present at this node; see Figure 3c). This is done for all spacer positions in the MSAA and all PSIO-contradicting events along the tree.
3. Compute the maximum joint likelihood reconstruction with the added fixed ancestral states that help to resolve the contradictions. Note that the newly reconstructed insertion events may not be exactly at the fixed nodes, but can be reconstructed even closer to the root.

Then go to step 1. The added fixed ancestral states are retained through all repetitions of this process.

For more details about this process see Supplementary Note 2.

#### Classification of spacer duplication candidates

The computed MSAA allows us to identify spacer candidates that are likely the result of spacer duplication, rearrangement, reinsertions, or independent acquisition events, since they are present in multiple columns of the alignment. Each identified candidate is relabeled such that each spacer/column in the MSAA is unique. For example, in Figure 3, spacer 3 has a set of two duplication candidates *{*3A, 3B*}*.

All of these event types share the characteristic of not conforming to the rules of polarized acquisition. As a result, they have the potential to cause misleading conflicts within the PSIO, e.g. when both versions of a duplicated spacer are included in the PSIO. To overcome this problem, we adopt the following approach:

Since each duplication candidate found in the MSAA is uniquely labeled, they are treated as independent spacers in the guide reconstruction. In the refinement step, only the candidate spacer that was acquired the earliest in the guide reconstruction remains in the PSIO and can produce conflicts with the PSIO in the guide reconstructions. Any conflicts to the PSIO arising from other duplication candidates are ignored at this stage. In Figure 3, spacer 3B is found to be the “oldest” candidate and therefore remains in the PSIO, while 3A (dashed) is not considered to be part of the PSIO.

After refinement, candidates (excluding the “oldest” candidate), or rather their insertion events, are classified according to the reconstructed evolutionary events of all candidates. We distinguish between rearrangements, reacquisitions, duplications, independent acquisitions, and other more complex types of duplicate events. In Figure 3, the insertion of 3A on *s*_0_ is classified as a duplication event, since 3B is already present at the root and 3A is in the middle of the array.

For a detailed description and illustration of the classification procedure and the different classes see Supplementary Note 2. More than 96% of the spacers are in line with polarized insertions and deletions (Table 2), justifying this classification step outside the scope of the model-based reconstructions.

#### Parameter estimation

We estimate the spacer deletion rate *ρ* and the mean block length of spacer deletions *α* by numerically maximizing the likelihood given by the independent deletion model or blockwise deletion model for the *reconstructed* spacer deletions.

For a group of spacer arrays, let *𝒯* be the corresponding phylogenetic tree and *N*_*b*_ the number of spacers present at the parent node of branch *b ∈ 𝒯*. Also, let *K*_*b*_ describe the set of reconstructed spacers that are deleted along branch *b* such that |*K*_*b*_| is the total number of spacers deleted at branch *b*. Moreover, let *K* := *{K*_*b*_ | *b ∈ 𝒯 }* and *N* := *{N*_*b*_ | *b ∈ 𝒯 }* be the collections of these sets for all branches in the tree.

We can compute the likelihood *L*_B_(*b, N*_*b*_, *K*_*b*_, *ρ, α*) of deletions *K*_*b*_ to occur along branch *b*, while retaining *N*_*b*_ *−* |*K*_*b*_| spacers, under the block deletion model with deletion rate *ρ* and mean block length *α*. Then the total likelihood of a group under the BDM is given by

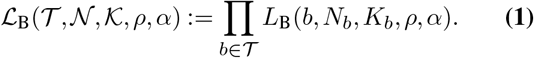

To compute the likelihood under the IDM we simply set *α* = 1, i.e. *ℒ* _I_(*𝒯*, *𝒩*, *𝒦, ρ*) := *ℒ* _B_(*𝒯*, *𝒩*, *𝒦, ρ*, 1). The (analytic) computation of the likelihood function of the block deletion model *L*_B_ is not trivial. The main issue to solve is that a connected component of consecutive spacers that have been lost along a branch of the phylogeny can arise from a combination of multiple block deletion events of different lengths. The two approaches we use to calculate the combined likelihood are described in detail in Supplementary Note 4. As the results are based on the branch lengths of the phylogenetic tree, the estimated deletion rate 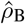is proportional to the time scale of this tree. In the core genome-based trees used here, the branch length is scaled as the expected number of substitutions per site. In this scaling, an estimated spacer deletion rate of 374 indicates that a spacer is deleted 374 times more frequently than a single site mutates.

#### Bias correction for unobserved extinct spacers

Spacers that existed at some time in the past can get deleted such that these spacers are not observed in any of the sampled spacer arrays. Consequently, deletion events related to such spacers cannot be reconstructed. This results in an underestimation of the deletion rate *ρ* in both the IDM and the BDM. Therefore, we correct the likelihood for such unobserved spacers by conditioning on maintaining the spacers in at least one of the samples that are descendants of the individual where the spacer has been acquired, similar to an approach introduced by Joseph Felsenstein in a different setting [49].

For the BDM the exact computation of this unobserved likelihood is hard due to the possibility of arbitrarily long deletions originating from unobserved spacers. Thus, we approximate the effect of the conditioned likelihood in the BDM using the correction factor from the IDM. For a precise description of the computation of the correction factor see Supplementary Note 4D. The effects of unobserved spacers on parameter estimates are significant (Supplementary Figure S8), but our bias correction can effectively address this issue.

Moreover, the maximum likelihood estimator of a geometrically distributed random variable is known to be biased, especially for a low number of observations. Therefore, in addition to the IDM-based bias correction for unobserved spacers, we use a standard bias correction for the estimated mean spacer deletion block length *α* after the numerical estimation step; see Supplementary Note 4D.

The improvement of parameter estimates by the combination of both bias corrections is evident when comparing Figure 5 and Supplementary Figure S9.

**Fig. 5.**
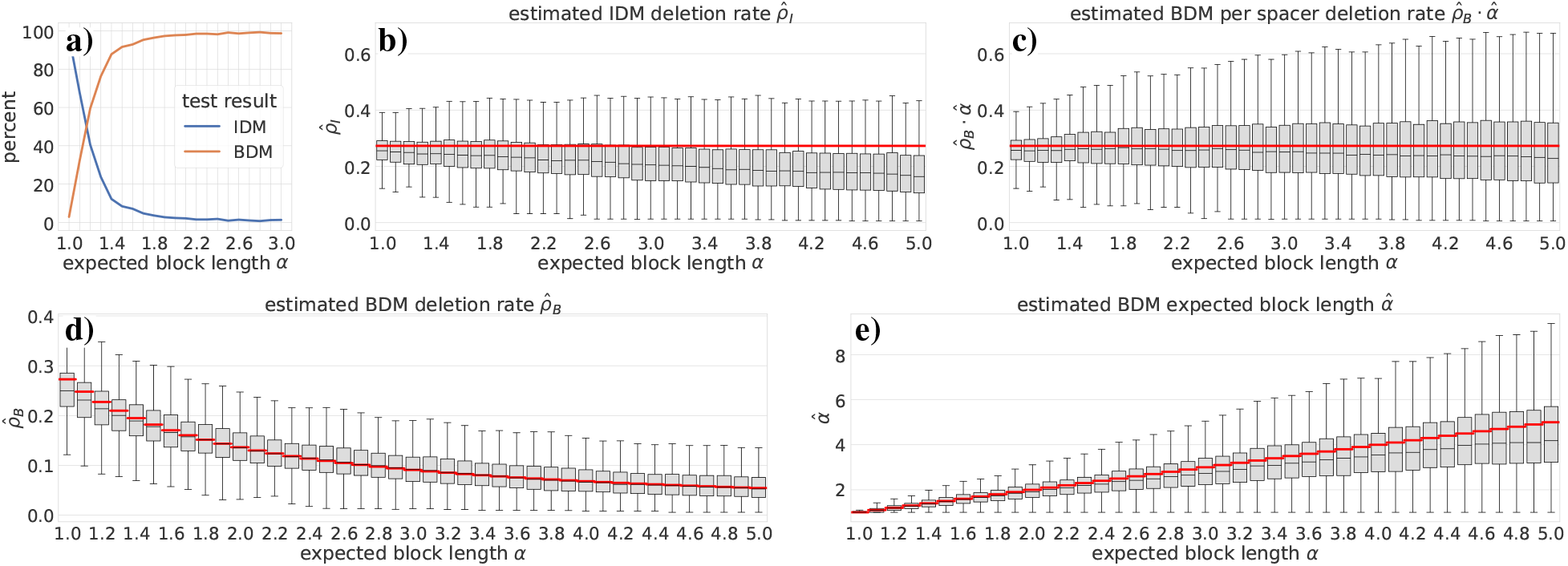
Parameter estimation and LRT-performance on simulated data. We evaluated *SpacerPlacer* on simulated spacer arrays. The simulations are based on random coalescent trees for different average lengths of blocks of jointly deleted spacers (*α*), where the deletion initiation rate *ρ*_B_ in the BDM was always chosen such that the per spacer deletion rate *ρ*_B_ *· α* remains constant for different choices of *α*. True parameters are shown as red lines. *SpacerPlacer* was then used to reconstruct the ancestral spacer arrays and estimate the corresponding parameters (using the simulated coalescent trees and including both bias corrections), shown as boxplots here. **a)** The percentages of likelihood ratio tests (LRTs) that indicate a BDM (orange line) or do not reject the IDM (blue line) are shown. For *α >* 1.6 the likelihood ratio test in *SpacerPlacer* can correctly recognize the BDM used in the simulation (ground truth). **b)** Estimates of the spacer deletion rate *ρ*_I_ based on the IDM. One can clearly see that the accuracy of the estimated per spacer deletion rate (difference to red line) drops with the expected block length *α* used in the simulations. This indicates that the IDM is not sufficient to predict the accurate average deletion rate per spacer when deletions occur in larger blocks. **c)** Estimates based on the BDM for the per spacer deletion rate *ρ*_B_ *· α*. As expected, parameter estimation based on the BDM performs better than the estimation based on the IDM in part b). Furthermore, the performance is strong for all values of *α*, only decreasing for high values of *α*. Overall *SpacerPlacer* estimates the per spacer deletion rate *ρ*_B_ *· α* well in the median. **d)+e)** Estimates of BDM deletion rate (bottom left) and BDM block lenght *α* (bottom right) separately. The result from part c) can be decomposed into the estimates of *ρ*_B_ (bottom left) and *α* (bottom right). As their product is fixed, the underlying *ρ*_B_ used in the simulations (red curve) decreases with growing *α*. This indicates that not only the per spacer deletion rate, but also the deletion initiation rate and the deletion length are well estimated in the median. There is a trade-off between *ρ*_B_ and *α* since both parameters are estimated simultaneously.

#### Likelihood ratio test

Based on the likelihoods computed in the previous section, we formulate a maximum likelihood ratio test to investigate within *SpacerPlacer* whether spacers are (likely) deleted in blocks or individually.

Here, a likelihood ratio test can be used, as the independent deletion model is nested in the block deletion model (setting *α* = 1). We formulate the hypotheses which correspond to the independent and block deletion model

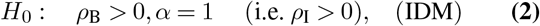

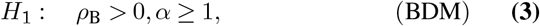

where the test statistic is given by

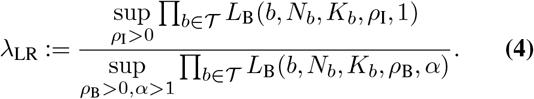

If *λ*_LR_ is significantly small, we dismiss *H*_0_ on *𝒯* and accept *H*_1_, i.e. the block deletion model fits the data better and is more likely. Otherwise, we cannot dismiss *H*_0_, i.e. both models explain the data reasonably well. The test is illustrated in Figure 4.

See Supplementary Note 4 for a more detailed description of the likelihood calculation and the likelihood ratio test.

#### Parametric models enable accurate estimation of the rates and lengths of spacer deletions

We evaluated the performance of *SpacerPlacer* using simulated data (Figure 5). *SpacerPlacer* performs well in estimating the (median) deletion rates in simulations and counteracts the bias of likelihood models towards too few events by incorporating the information from the PSIO, indicating a well-performing reconstruction algorithm (Figure 5 (b, *α* = 1), (d), and (e)). The likelihood ratio test reliably distinguishes independent individual spacer deletions compared to joint deletion of adjacent spacers, as soon as the average length of jointly deleted blocks of spacers exceeds 1.6 in the simulations (Figure 5(a)).

In simulations where spacer deletions occur in blocks, assuming an IDM model results in an underestimation of the per spacer deletion rate (Figure 5(b)). Thus, it is important to incorporate block deletions into the parametric model to correctly estimate the frequency and length of spacer deletions. When the BDM is set as the underlying model, *SpacerPlacer* is able to accurately estimate the per spacer deletion rate and the parameters *ρ*_B_, *α* individually (Figure 5(d), and (e)). The estimation quality only degrades for very high *α* when deletion events that collapse (almost) the entire array occur more frequently. If the effective rate of deletion per spacer, calculated by taking the product from *ρ*_B_ and *α*, is considered (Figure 5(c)), the IDM and BDM estimates are more comparable. Here, the tendency to underestimate the deletion rate when using the IDM becomes clear (see Figure 5(b)). *SpacerPlacer* showed similar performance for simulations based on the parameter estimates and core-genome trees from the CRISPRCasdb dataset (Supplementary Figure S10).

The estimation of parameters in the BDM is non-canonical due to the dependencies created by jointly lost neighboring spacers (Supplementary Note 4). In addition, we need to use bias corrections for both *ρ*_B_, *α*. BDM-based estimates without bias corrections tend to underestimate *ρ*_B_ and *α*, as can be seen in Supplementary Figure S9. Due to the typically short length of spacer arrays, the variance of the estimated rates and block lengths is high for individual groups, suggesting that the estimated parameters are more reliable in well-curated, large groups and when averaging over multiple CRISPR array groups.

### Analyzing the patterns of spacer array evolution across numerous bacterial genomes

The benefits of our modeling framework are best highlighted in the context of the dynamics of spacer deletions and the postulate that such deletions are caused by replication slippage. Deletions that arise from repeat misalignment introduce three effects that can be detected in the inferred parameters and reconstructions based on the BDM. First, near both ends of the array, the observed deletion frequencies will be reduced because repeats at the boundary of the array have a lower number of nearby repeats with which they can align for a deletion event. We call this the “array boundary effect” on the deletion frequency. Second, misalignments spanning multiple repeat-spacer units can result in patterns where the simultaneous deletion of multiple adjacent spacers can be observed. In this case, the likelihood ratio test should favor the BDM and indicate how frequent and long such deletion blocks are. Third, deletions by repeat misalignment are, a priori, independent of the Cas genes, i.e. the Cas type.

#### Distribution of deletions along CRISPR arrays aligns with deletions caused by replication misalignments between repeats

To investigate the distribution of deletions according to their position in the spacer array, we compared *SpacerPlacer* reconstructions for two different simulation scenarios (Figure 6). We either assumed that spacer losses occur evenly distributed along the array without the need to pair different repeats or, in the second scenario, that spacers at the edges of the array are less often part of block deletions according to the lower number of neighbor spacers and repeats that could mismatch with them. To avoid counter-intuitive binning effects, we only show groups with an average array length between 15 and 25 in Figure 6. We provide similar results for further array lengths, a differently scaled visualization, and additional details in Supplementary Figures S11, S12 and S13.

**Fig. 6.**
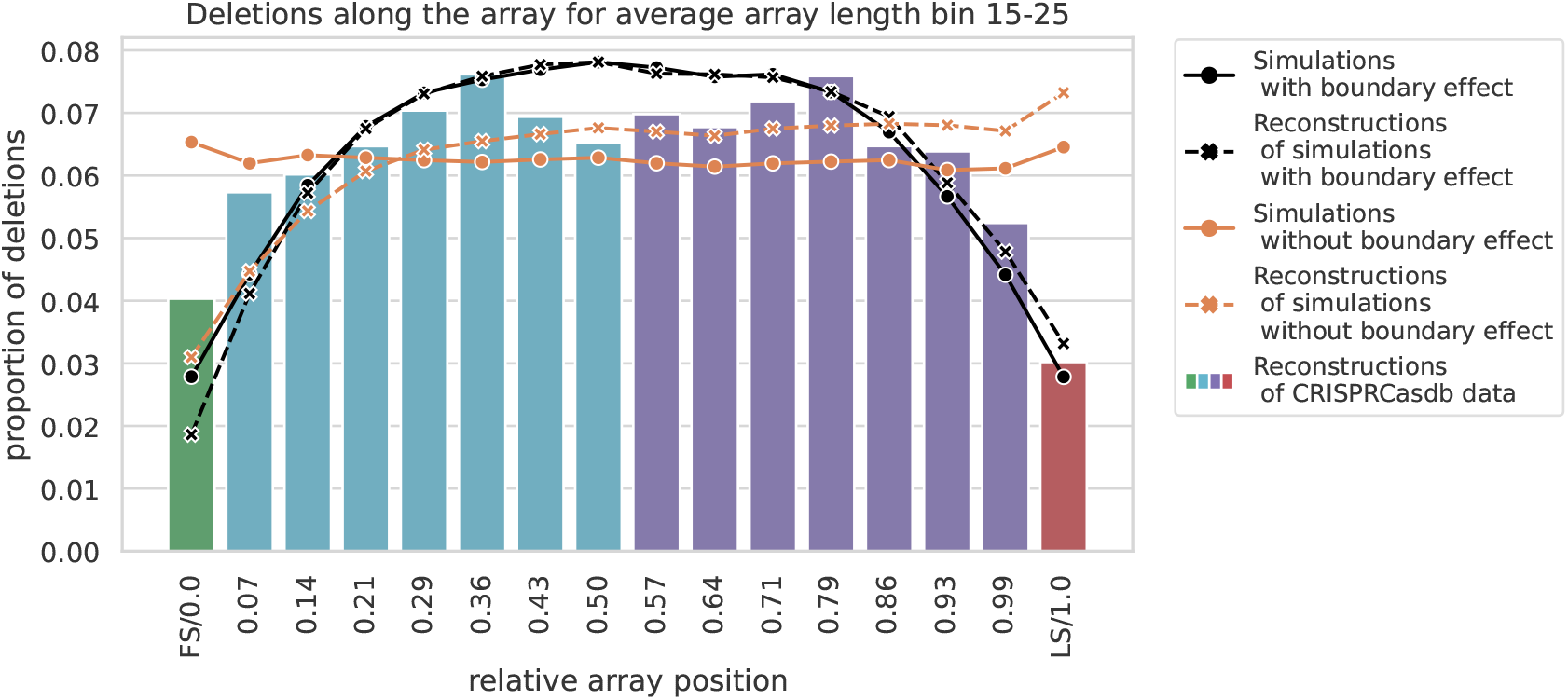
Proportions of deletions with respect to the position along the array. The solid orange line shows the proportion of deletions for simulations under an artificial block deletion model, where deletions of internal spacers and spacers at the ends of the array occur at the same frequency. In contrast, the solid black line shows the proportion of deletions for simulations under the more natural block deletion model where block deletions cannot exceed the array boundary as nearby repeats need to align. Here, deletions are less likely to occur towards both ends. The dashed lines show the corresponding values based on the events reconstructed by *SpacerPlacer* in colors corresponding to the respective models. In both cases *SpacerPlacer* is able to reconstruct the deletion distribution of the respective models, except near the leader end of the array. The underlying histogram shows the distribution of deletions in reconstructions based on CRISPRCasdb data which is similar to the deletion model with a boundary effect (solid and dashed black lines). But there are more reconstructed deletions at the first spacer position than at the last spacer position, contrasting the pattern we would expect based on the neutral simulations. For each category, we bin all deletion events according to their relative position compared to the reconstructed lengths of the array. Each point or bar corresponds to a bin, with the label on the x-axis indicating the upper limit of the bin. The y-axis shows the proportion of deletions falling within the respective bin. The first spacer (FS/0.0) and the last spacer (LS/1.0) are placed in their own bins (shown in green/red).

The PSIO is particularly advantageous for the reconstruction of older spacers, as more of the younger spacers can provide information about the correct insertion time. In both simulation scenarios, the accuracy of ancestral reconstruction towards the leader end, where new spacers are acquired, is thus lower than towards the trailer end of the array (Figure 6). Therefore, we expect that reconstruction at the leader ends for CRISPRCasdb data are similarly impacted.

The frequency of reconstructed spacer deletions based on CRISPRCasdb data shows a fall-off towards both boundaries of the array with a roughly uniform distribution in the middle of the array (Figure 6). This is different from what we would expect from a distribution that is directly dependent on the length of the array, such as the fragment deletion model proposed by [36] or models that favor deletions on one side of the array [8, 30]. Interestingly, a uniform deletion rate along the array was also found to be a representative model for spacer deletions in a related simulation study [50]. The uniform distribution in the middle combined with the drop towards both ends of the array is consistent with a pairwise repeat-dependent deletion process [12] with geometrically distributed deletion lengths.

#### Blockwise deletions of neighboring spacers are frequent

Several theories and mechanisms have been proposed to explain the distribution of joint deletions [12, 36, 51, 52]. Until now, however, statistical tests have not been able to determine the most likely scenario, as the deletion patterns observed in different CRISPR array datasets are consistent with different models for spacer deletions. In particular, both the IDM and BDM contain losses of adjacent spacers that can arise from combinations of multiple overlapping deletions; only their frequency and distribution change.

To determine the mode of spacer deletions in CRISPR arrays, we applied the likelihood ratio test described in a previous section to the datasets extracted from CRISPRCasdb and reconstructed the ancestral CRISPR arrays to investigate the properties of spacer array evolution.

We found that in 76% of the CRISPR array groups, the likelihood ratio test rejected the null hypothesis, i.e. the IDM, in favor of the BDM. For a breakdown of the test results with respect to Cas type; see Supplementary Figure S14. Although the IDM cannot be rejected in the remaining cases, this indicates that, overall, the block deletion model describes the natural process of spacer deletions significantly better.

We inferred the distribution of the lengths of the joint deletions in our reconstructions based on CRISPRCasdb data (Supplementary Figure S15). The inferred empirical distribution is consistent with a geometric distribution for smaller lengths but, notably, does have a heavier tail. Thus, we hypothesize that the deletion process may, to a lesser extent, involve additional mechanisms, such as fragment deletions and the loss of entire CRISPR arrays.

For the expected block length parameter *α* the median of all estimated parameters based on the CRISPRCasdb datasets is *α ≈* 2.73. This is consistent with our statistical analysis of the block deletion lengths observed by Lopez-Sanchez et al. [38]. The length of the spacer blocks jointly deleted in their experiment fits a geometric distribution (p-value = 2.76 *·* 10*−*5) and has a mean deletion length of *α* = 3.32.

Therefore, in all subsequent analyzes, we will only present the results derived from the more accurate BDM, unless otherwise indicated.

#### Spacer deletions are frequent and Cas type-independent

The estimated spacer deletion rates and lengths are comparable for the different Cas types investigated here (Figure 7), indicating that, no Cas-dependent molecular mechanism influences the deletion of spacers. We find similar results with respect to genus (Supplementary Figure S16).

**Fig. 7.**
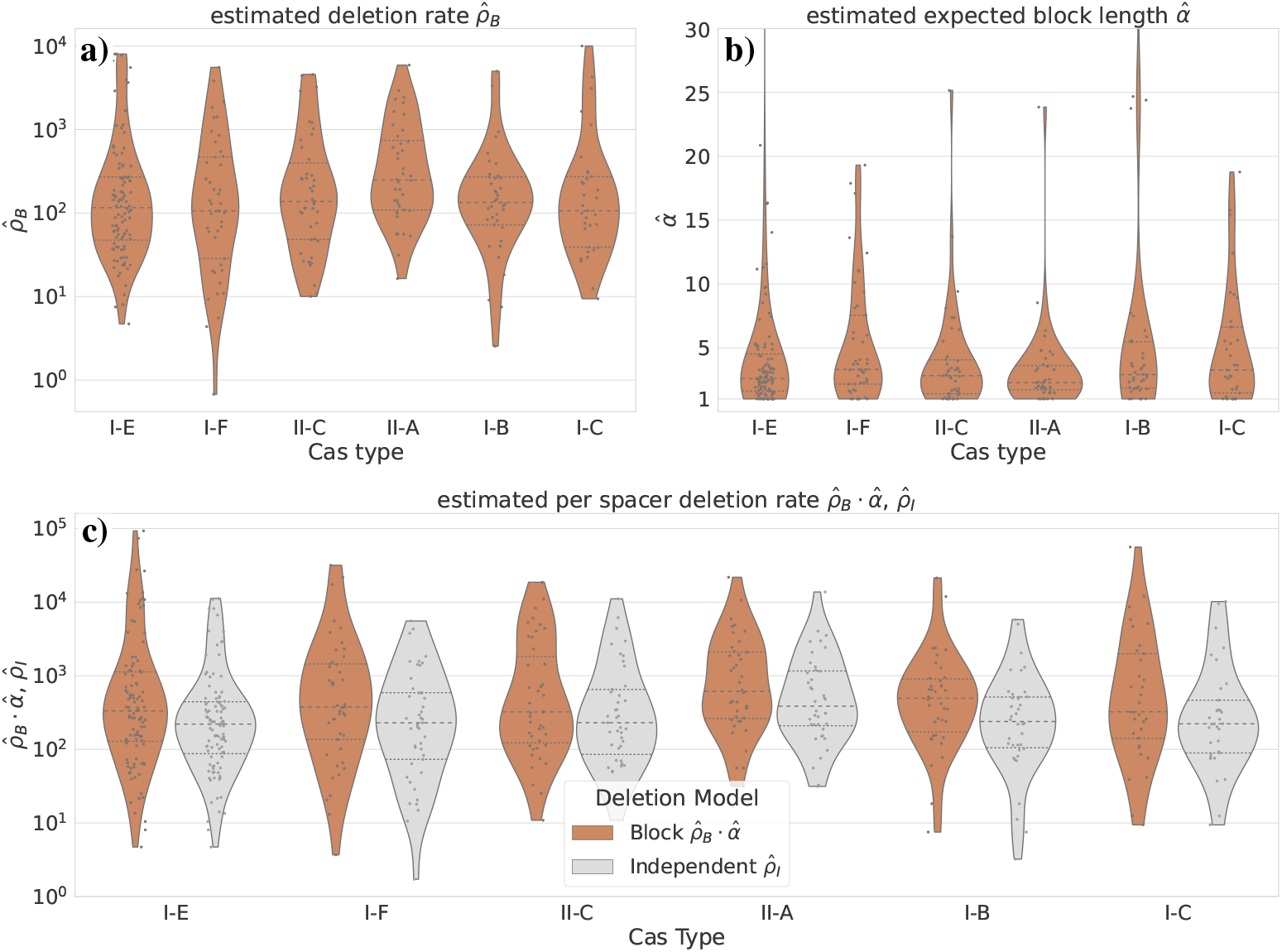
Parameter estimates on CRISPRCasdb data. The estimated parameters for **a)** the deletion initiation rate, and **b)** the expected length of deletion events in a block deletion model, as well as **c)** the resulting per spacer deletion rate for different Cas types are shown. There are no significant differences between the Cas types. The variation of individual estimates is high, but the median estimates of the per-spacer deletion rate are consistently similar. For comparison, we also show the estimates, if the underlying model is the less accurate IDM (light gray) in (c). Although IDM-based estimates correlate with BDM-based results, the same bias as in the simulations can be observed. See Table 1 for the global mean and median parameters.

We can estimate the evolutionary turnover speed of CRISPR arrays because the core genome trees used for the estimates are time-scaled based on the average per site mutation rate. In the BDM, we estimate the deletion initiation rate *ρ*_*B*_ and the mean block length *α* of deletions. From these we can calculate the average deletion rate per spacer *ρ*_B_ *· α*. Global median and mean estimates are shown in Table 1. The median of the spacer deletion rate per spacer in the BDM is 374, indicating that a spacer is deleted at a rate 374 times higher than the rate at which individual nucleotide mutations occur. In particular, since spacers (and their repeats) are generally smaller than 100 bp in length, the deletion of spacer sites occurs at a faster rate than nucleotide mutations within the array. The loss of immunity by spacer deletion is therefore the greater risk for the host compared to mutations within the spacer.

**Table 1.**
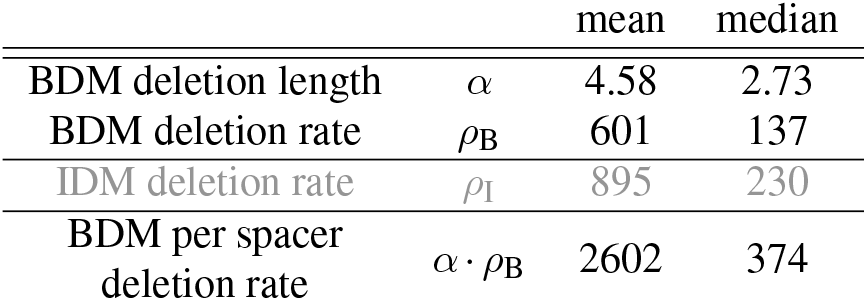
Mean and median parameter estimates in the independent and block deletion model. We estimated the parameters of the independent and block deletion model on CRISPRCasdb data across all groups with at least one reconstructed deletion; see also Figure 7. Top: Estimates for the BDM. In the BDM deletions are initiated at a rate *ρ*_B_ and cover a geometrically distributed number of spacers with mean *α*. Middle gray: Estimate of the deletion rate *ρ*_I_ of the IDM. In this model, all initiated deletions have length 1. Bottom: *α· ρ*_B_ is the approximate deletion rate per spacer in the BDM and roughly corresponds to *ρ*_I_ in the IDM.

We can also give an ad hoc estimate of the median insertion rate *θ ≈* 6522 by considering the following. As each individual spacer is lost at a mean rate of *ρ*_*B*_ *· α*, an overall spacer insertion rate of *n · ρ*_*B*_ *· α* will result in a balance that sustains an average array length of *n ≈* 17.4. Note that the insertion rate represents the total rate of acquisition of new spacers and thus is naturally significantly higher than the per spacer deletion rate.

This indirect rough estimate of the average insertion rate should be considered with extreme caution, especially for single groups, where the variance of the estimates is large (Supplementary Figure S17). As the frequency of spacer acquisitions depends on environmental fluctuations such as co-evolution with phages, we would expect the insertions to occur in bursts with varying rates, while the model assumes a constant rate. Thus, it remains unclear how an inferred average acquisition rate can be interpreted.

In contrast, spacer deletions can take place without the involvement of any foreign elements, which suggests that they can be better explained by a constant rate. However, together the estimates indicate that spacer turnover is fast.

#### Spacer duplications and rearrangements are rare

Spacers that can be explained solely by insertions and deletions were by far the most common case in the CRISPRCasdb datasets, but we also identified roughly 3% of all spacers that form loops in the PSIO and are only explainable by more complex event types, as shown in Supplementary Note 2, Figure S1. Of these complex events, 86% are simple duplications, 5% are rearrangements, and only 1% new insertions of the same spacer (Table 2). This aligns with the finding of Kupczok et al. that recombination between CRISPR arrays is rare [51].

**Table 2.** Number and proportions of duplication events. The total number of spacers of the CRISPRCasdb dataset is divided into spacers that are compatible with polarised insertions and deletions only, and spacers that are duplication candidates according to the MSAA and the PSIO. Duplication candidates are further classified into reconstructed complex event types.

|  | count | proportion |
| --- | --- | --- |
| total number of spacers | 24,090 |  |
| PSIO-compatible spacers | 23,285 | 0.967 |
| duplication candidates | 805 | 0.033 |

**Table 2. Number and proportions of duplication events.**
|  | count | prop. of candidates |
| --- | --- | --- |
| duplications | 695 | 0.863 |
| rearrangements | 43 | 0.053 |
| reacquisitions | 8 | 0.010 |
| independent insertions | 7 | 0.009 |
| other events | 52 | 0.065 |

#### No evidence of age-dependent selection along the spacer array, except possibly at the first and last positions

If selection pressures would differ along the array, e.g. if more anciently acquired spacers would provide a higher or lower fitness benefit than more recently acquired spacers, one would expect that the observed deletion rate along the array correlates with this selective advantage. It is important to distinguish between internal spacers and spacers that are close to the first or last position. For the later, we expect a strong additional fitness-independent influence of the underlying repeat-dependent blockwise deletion process which causes lower deletion rates towards the array boundary. When looking at the relative position of reconstructed deletions, the frequency of deletions are overall in line with the boundary effect in the BDM (Figure 6). At least for spacers that are not among the very first or last positions, the absence of a clear sign of a selective gradient indicates that the selective pressures to maintain a particular beneficial spacer and the fitness costs to keep a useless spacer are either constant over time or small.

For the first (and last) spacers, the task of inferring the selection strength is particularly challenging, as the bias created by the reconstruction is not symmetric and can obscure the analysis (Figure 6). To take this reconstruction effect into account, we compared the frequencies of reconstructed CRISPRCasdb deletions to reconstructions from simulated spacer arrays. Of the perfectly symmetric simulated deletions *SpacerPlacer* reconstructs fewer deletions at the first position compared to the last position. In contrast, the CRISPRCasdb data show more reconstructed deletions of the first two spacers compared to the last two spacers (Figure 8). This suggests that the spacer deletion rate is higher towards the leader end. In the discussion, we describe potential selective benefits of losing recently acquired spacers and alternative explanations that could cause this difference.

**Fig. 8.**
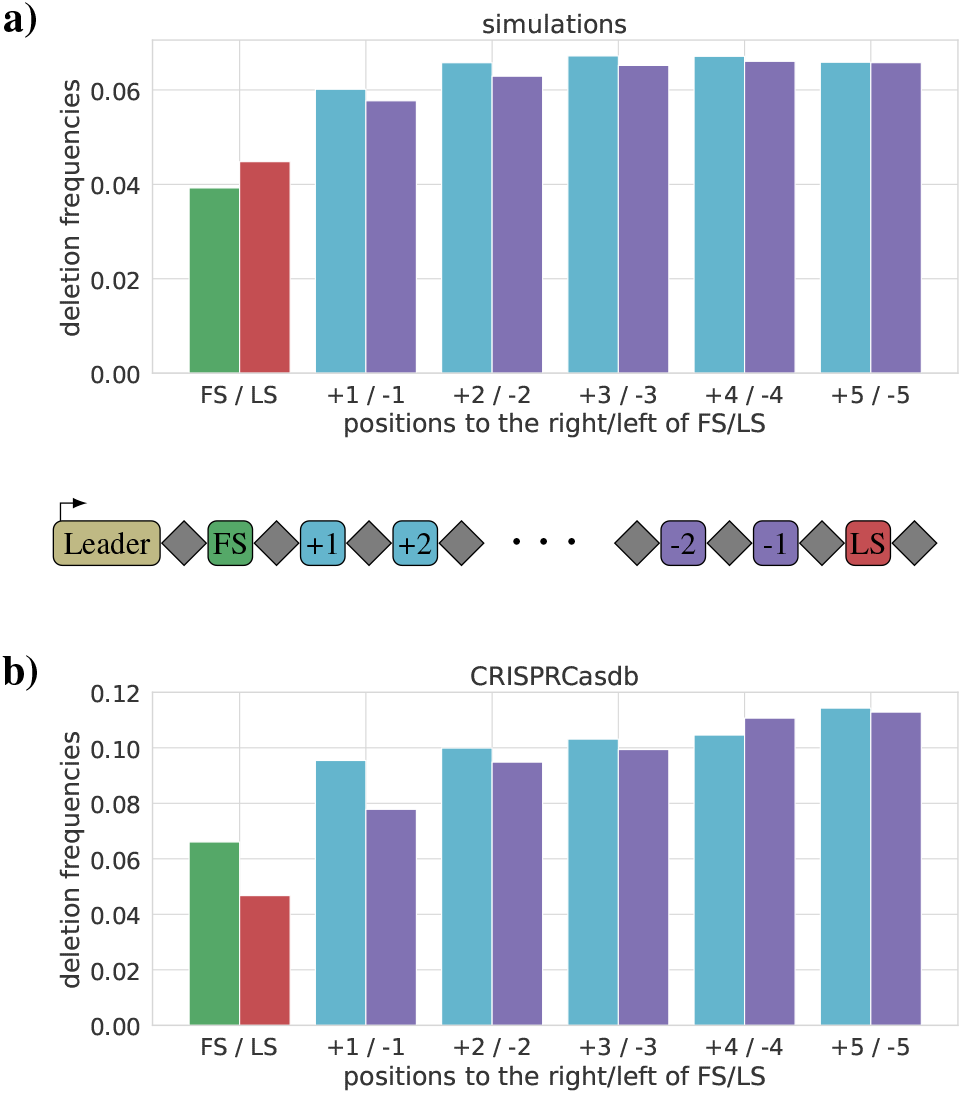
Deletion frequencies with respect to the proximity to the array boundary. We show the deletion frequencies for absolute positions next to the first and last spacer. **a)** shows the reconstructed deletion frequencies based on simulations with array boundary effect. The first spacer (FS) appears to be deleted less often than the last spacer (LS), although the simulation is perfectly symmetric. But, just as in Figure 6, the reconstruction is more likely to underestimate the frequency of FS deletions. **b)** shows the deletion frequencies for reconstructions based on CRISPRCasdb data. Here, the pattern is reversed, and the LS is less frequently deleted than the first spacer. Comparing the FS and LS deletion frequencies between a) and b) suggests that compared to a symmetric model, either the LS is deleted less frequently, the FS is deleted more frequently, or both.

## Discussion

### Comparison to other models of spacer array evolution

Multiple modeling approaches explored the coevolutionary dynamics between phages and bacteria [30, 50, 53] and the length of spacer arrays [54, 55]. In addition, some stochastic models have been specifically designed for CRISPR spacer array evolution [36, 37]. Kupczok and Bollback suggested a spacer deletion model in which any fragment of neighboring spacers of the array is equally likely to be deleted. Here we propose geometrically distributed block deletion lengths which fit well with observed deletion lengths in CRISPRCasdb and experiments [38] and allow efficient likelihood computations and parameter estimations.

Collins and Whitaker introduced a reconstruction tool based on maximum parsimony principles [8] based on empirically determined scores for different events in CRISPR array evolution. In contrast, we rely on explicit evolutionary models that can offer insight beyond ancestral reconstruction through parameters that correspond to molecular mechanisms and evolutionary processes.

A detailed discussion and overview of the models used in related studies and different steps of our pipeline, including our LRT workflow, is shown in Supplementary Note S5.

### Using the partial spacer insertion order is crucial to reconstruct ancestral spacer deletions

Inferring the dynamics of spacer array evolution is challenging, as arrays are typically short and provide very limited information. Furthermore, the acquisition of new spacers correlates with environmental conditions, such as the abundance of phages. In contrast, spacer deletions are less dependent on the environment, and thus are more likely to follow a molecular clock. Fortunately, the order of spacers provides additional information and repeated acquisitions of the same spacer are extremely rare and thus can be safely ignored. These two properties form the basis for the reconstruction approach. *SpacerPlacer* first reconstructs the most likely acquisition point(s) and the resulting deletions along the tree independently for each spacer. And afterwards refines these according to the PSIO.

Enforcing the chronological order of spacer insertion during the refinement step based on the PSIO counteracts the tendency of likelihood models to explain patterns by fewer (deletion) events. This allows *SpacerPlacer* to reconstruct ancestral spacer arrays with greater precision and improves rate estimates. In the CRISPRCasdb data, more than a quarter of the spacers (6,773 out of 24,090) are affected, i.e. their acquisition events are relocated during the refinement procedure. This results in a higher number of reconstructed spacer deletions that are subsequently used by *SpacerPlacer* to estimate the maximum likelihood parameters in the more complex evolutionary model with blockwise spacer deletions.

### Spacer deletion patterns are independent of the Cas type

Our results indicate that joint deletions are a common phenomenon in CRISPR array evolution and show no significant differences in their deletion rates or lengths between Cas types I-B, I-C, I-E, I-F, II-A, and II-C based on CRIPSRCasdb data. This suggests that spacer deletions are not controlled or regulated by CRISPR-specific molecular mechanisms, but rather occur in a random fashion. Quite contrary to this view, frequent spacer deletion and renewal of spacer composition is essential for the effectiveness of the system [54] and could, in principle, have fostered the emergence of a dynamic deletion system similar to primed adaptation. Interestingly, deletion frequencies can be subject to selection indirectly when multiple CRISPR arrays with different lengths, and therefore due to the boundary effect also different per spacer deletion rates, coexist [56, 57].

Although we did not find significant differences here, the parametric models behind *SpacerPlacer* will allow future studies on a larger CRISPR database to identify potential differences in deletion patterns.

The parameters estimated in groups of spacer arrays from CRISPRCasdb reveal two interesting characteristics of spacer array evolution. First, the deletion rate per spacer is 374 times higher than the mutation rate per site, highlighting that immunity loss via spacer deletion acts on a much faster time scale compared to mutations within spacers.

Second, in line with deletions that arise from repeat misalignments, blocks of jointly deleted spacers occur frequently, and their length distribution is consistent with a geometric distribution with mean 2.7 that is interrupted if the block length exceeds the spacer array. Nonetheless, single-spacer deletions remain the most common event (37% for *α* = 2.7).

### Reconstructed duplications shed light on the mechanisms of spacer deletions

Our analysis of CRISPRCasdb spacer evolution showed that complex events, including duplications, rearrangements, and reacquisitions, are rare and dominated by duplication events. Only 3% of the considered spacers have been reconstructed as duplicated, and 171 out of 333 CRISPR array groups do not include a potential duplication. The number of observed duplications is much lower than we would expect from the experiment in [38] where roughly 10% of the arrays had a duplication. There are two explanations: 1) Our model only provides a lower bound for such events, and 2) duplicated spacers provide an additional possibility for replication slippage since the two identical spacers can also misalign. It is conceivable that duplicated spacers are thus lost at a higher rate than nonduplicated spacers, leading also to a lower rate of detected duplications. This observation adds an interesting aspect to the postulate that most spacer deletions occur due to replication misalignment (“slippage”) of the repeats [12, 40].

### Deciphering selection pressures from deletion patterns in spacer arrays

One of the main questions motivating this work was whether spacers at different positions in the spacer array are under different selection pressures. The parametric model-based approach allows us not only to investigate the likelihood of evolutionary forces and molecular mechanisms at play, but also to reveal hints at selective pressures acting on the presence and absence of spacers at different positions in the array.

It is important to distinguish the position along the reconstructed ancestral array and the current position among the observed sample arrays. When looking at the deletions in the spacer arrays of a sample, these seem to appear predominantly at the distal end. However, deletions of older spacers are naturally observed more often in such alignments, since they existed for a longer time during which deletions can occur compared to more recently acquired spacers. At the time when the deletions occurred, their position was closer to the leader end. This observation bias has led to the perception within the scientific community that deletions occur more frequently among shared spacers at the end of the array, rendering them less beneficial than more recently acquired spacers, which seem to be less often deleted.

Our results indicate that this impression is a misinterpretation of the accumulation of observed deletions towards the distal end in alignments of current spacer arrays. Here, we correct this misleading impression by instead considering the reconstructed deletion positions in ancestral spacer arrays. We observe no clear sign of selection acting differently on deletion frequencies along the reconstructed arrays, at least for spacers that are not among the first or last positions. While we by no means propose that CRISPR arrays evolve neutrally, our results hint at either overall weak selection pressures to delete specific spacers or an equally strong selection along the array to maintain spacers.

We found that the drop in deletion frequency towards the edge of the array is particularly pronounced for the last spacer compared to the first spacer. This difference could be caused by the often observed excessive mutation of the last repeat [10], which renders them less likely to participate in deletion events based on misaligned repeats.

Vice versa, we can argue that the difference between the last and first spacers arises because of selective effects at the leader end. As we can only reconstruct deletions from individuals whose offspring survived, this could indicate that many recently acquired spacers are not beneficial in the long run and can be deleted again quickly. Another selective advantage of deleting spacers at the leader end could arise due to the expression levels along the spacer array. Modifying the composition of spacers at the first most expressed positions might be particularly beneficial and could lead to the observed increased deletion rates at the first spacer positions.

Interestingly, our findings are in line with recent observations by López-Beltrán et al. [25]. They showed that the environmental abundance of the phages and plasmids targeted by spacers varies along the spacer array. They found that at the first positions, the targeted phages are present in the environment, whereas the targets of the spacers directly following are less abundant. This indicates a lower selection pressure to maintain recently acquired spacers. Toward the trailer end they observed that the spacers correspond to more abundant but different phages and plasmids, which fits well with the reconstructed deletion frequencies here.

### Dataset quality and accuracy of the underlying tree affect spacer array reconstructions

Provided there is sufficient overlap to form a PSIO, including incomplete arrays should not significantly affect the reconstruction. However, the impact of an incorrect tree on the reconstructed ancestral arrays could theoretically be large. Misplaced tips can cause chain reactions during the refinement step, such that more and more spacer insertions are wrongly moved toward the root. However, closely related sister arrays are rarely placed far apart and a minor misplacement will cause limited damage. In addition, such effects are apparent and likely to be observed by a knowledgeable user. Comparing the spacer-array based tree from *SpacerPlacer* with the core phylogeny can be a good starting point for this. Such a comparison can also help identify whether multiple spacers within a sample are artifacts of the CRISPR detection process, such as a mistakenly split spacer array, or related coexisting CRISPR arrays [57]. The user can then resolve these problems by adjusting the tree, excluding arrays, removing specific spacers, or adjusting the parameters of *SpacerPlacer*.

Lastly, since the number of deletion events is typically small, parameter estimates based on a single dataset have a high variance. This is readily apparent in the estimates on CRISPRCasdb data (Figure 7) and simulations (Supplementary Figure S10). In general, we recommend trusting parameter estimates only in the median over a large number of arrays and groups using properly time-scaled trees.

### Applications and current limitations

*SpacerPlacer* is a rapid and versatile tool for studying CRISPR arrays and systems, and we expect it to provide valuable insights in a variety of different application scenarios.

For example, we inferred that the deletion rate for CRISPR spacers, *ρ*_*B*_ *· α* = 374, is two orders of magnitude higher than the per-site mutation rate. This evolutionary speed and the robustness of complete spacers with regard to sequencing errors make CRISPR arrays a valuable fine-scale typing tool [19], as has already been demonstrated for multiple species [17, 20, 58, 59].

When tracking the evolution of clinically relevant samples and investigating outbreaks, it can in addition to CRISPR typing also be beneficial to reconstruct their evolutionary relationship. While standard multilocus sequence typing (MLST) is often insufficient to reconstruct the evolutionary history of closely related samples, CRISPR arrays are rich in phylogenetic information due to the PSIO. This concept was also discussed by Tomida et al. [14], who applied standard phylogenetic reconstruction tools on CRISPR arrays, which provided a higher resolution than the trees generated using MLST. The currently implemented block deletion model based distance in *SpacerPlacer* already solves some of the issues found in the CRISPR based trees of Tomida et al. (Supplementary Note 6A), and we anticipate that incorporating more advanced tree inference methods will further improve the results.

As a rough estimate of the discriminatory power of spacer arrays consider the following estimate. The number of deletions is proportional to the per spacer deltion rate times the length of the MSAA *n* (*≈* 38 in the CRISPRCasdb dataset).

Then to achieve a similar discriminatory power, a set of genes used for MLST needs to be of combined sequence length *n·ρ*_B_*α ≈* 38*·*374 *≈* 14200. Thus, spacer arrays have a roughly three to four times higher discriminatory power than MLST, where, typically around 3500 bp are considered [60]. Clearly, core-genome MLST and whole-genome based discrimination exceed this threshold and are thus more precise. However, spacer arrays have a high discriminatory power compared to their relative length and, unlike point mutations, provide temporal information encoded in the PSIO. Moreover, because of their repeat structure, they are easy to find and less susceptible to sequencing errors.

Monitoring bacterial samples and their co-evolution with phages based on CRISPR arrays is also of importance for bacteria-driven biotechnological and pharmaceutical production processes, such as bioreactors [23] or wastewater treatment plans [24] and other metagenomic time series [22– 24]. However, CRISPR arrays derived from short-read sequencing and metagenome assembled genomes are often incomplete. Naturally, the rate estimations will be obscured when using partial spacer arrays, but *SpacerPlacer* will still provide a partial reconstruction of ancestral spacer arrays. When complete arrays are available through long-read sequencing on metagenomic data or have been carefully reconstructed from short reads [59], *SpacerPlacer* benefits from the complete PSIO increasing the precision of ancestral array reconstruction and rate estimation. An example is Lam and Ye, who characterize CRISPR arrays and the corresponding PSIO graphs from a gut microbiome sequenced with long-read sequencing technologies [21]. In this situation, *SpacerPlacer* can infer the ancestral relationships for such datasets and reconstruct the corresponding spacer insertion and deletion events (Supplementary Note 6B).

In conclusion, *SpacerPlacer* is a new parametric ancestral reconstruction tool that enables in-depth investigations of CRISPR array evolution. Analyzing spacer arrays of closely related samples and the correspondence between spacers and invading elements within the reconstructions will allow us to track pathogens at a fine scale and shed light on the coevolution of phages and bacteria, the mechanisms of spacer acquisition and deletion itself, and the dynamic ecology of microbial communities and microbiomes.

## Data Availability

*SpacerPlacer* is available at https://github.com/fbaumdicker/SpacerPlacer. Example datasets are available in the github repository.

## ACKNOWLEDGEMENTS

We thank Shiraz Shah and Jaime Iranzo for their helpful comments and discussions. AF and FB are funded by the Deutsche Forschungsgemeinschaft (DFG, German Research Foundation) under the Priority Program - SPP 2141 - Project number Ba-5529/1-1. FB is funded by the Deutsche Forschungsgemeinschaft (DFG, German Research Foundation) under Germany’s Excellence Strategy – EXC number 2064/1 – Project number 390727645, and EXC 2124 – Project number 390838134.

This work was supported by German Research Foundation (DFG) [BA 2168/23-1 and BA 2168/23-2 SPP 2141]; Much more than Defence: the Multiple Functions and Facets of CRISPR–Cas; Baden-Wuerttemberg Ministry of Science, Research and Art; University of Freiburg.

This work was supported by the BMBF-funded de.NBI Cloud within the German Network for Bioinformatics Infrastructure (de.NBI) (031A532B, 031A533A, 031A533B, 031A534A, 031A535A, 031A537A, 031A537B, 031A537C, 031A537D, 031A538A). We acknowledge support by Open Access Publishing Fund of University of Tübingen.

