## Supplemental File for "*SpacerPlacer*: Ancestral reconstruction of CRISPR arrays reveals the evolutionary dynamics of spacer deletions"

[1]Cluster of Excellence “Controlling Microbes to Fight Infections”, Mathematical and Computational Population Genetics, University of Tübingen, Tübingen, Germany
[2]Institute for Bioinformatics and Medical Informatics (IBMI), University of Tübingen, Tübingen, Germany
[3]Bioinformatics group, Department of Computer Science, University of Freiburg, Freiburg, Germany
[4]Information and Computer Science Department, King Fahd University of Petroleum and Minerals (KFUPM), Dhahran 31261, Saudi Arabia
[5]Interdisciplinary Research Center for Intelligent Secure Systems (IRC-ISS), King Fahd University of Petroleum and Minerals (KFUPM), Dhahran 31261, Saudi Arabia
[6]Signalling Research Centres BIOSS and CIBSS, University of Freiburg, Freiburg, Germany
[*]shared first authorship

### Supplementary Note 1: Data extraction and simulations

**A. Data extraction from CRISPRCasdb and preprocessing.** The data was extracted from CRISPRCasdb [1] on June 30th 2022. A total of 143878 CRISPR spacer arrays were extracted that were grouped by their respective consensus repeats forming 53965 repeat groups. We refine the groups as follows (the remaining number of groups and arrays after each step is shown in brackets):

(a) For each repeat, the spacers were clustered, such that each cluster contains spacers which differ by a single mutation from one of the spacers in the same cluster. In this case, at least one spacer is only removed one point mutation from another spacer within the same cluster, others can be multiple mutations removed. Each cluster is labeled by a natural number. These clusters are generally small across all repeats within the dataset. Then each spacer within each array is replaced by the label of the respective cluster they belong to.

(b) We remove arrays with lower confidence ($< 3$) and only retain arrays with high confidence (4). (7352 groups with 32624 arrays)

(c) For arrays with uncertain orientation prediction, we checked if either the reverse complement of the group (repeat and spacers) has overlap with any group. If so, the groups were combined. (6525 groups with 32624 arrays)

(d) Arrays with no close (< 10000 bp distance) Cas-cassette are removed, unless only one Cas-cassette was found within the genome. The Cas type is assigned according to the closest Cas-cassette. (6122 groups with 29942 arrays)

(e) The groups are subdivided by genus and Cas-type. (7506 groups with 29942 arrays)

(f) The groups are split according to pairwise spacer overlap. For each spacer array there has to be at least one other array within the group with overlapping spacers. (12863 groups with 29942 arrays)

(g) We drop groups with less than 3 unique spacer arrays. (965 groups with 16853 arrays)

(h) Groups that exceed 1000 (non-unique) spacer arrays are dropped, since their impact on the parameter estimation is limited and their size leads to large increases of run time for the tree estimation. (4 groups with 7051 arrays are removed)

(i) Afterwards all spacer arrays are aligned with MAFFT and estimated trees based on their core-genome with IQ-TREE. Groups where no core-genome (annotations) are found are dropped.

We drop arrays that have multiple spacer arrays within a single group, since they can not be separated in the core-genome tree. We again remove groups that are reduced to below 3 arrays. (518 groups with 5934 arrays)

These 518 groups with 5934 arrays are fed into SpacerPlacer for our experiments on CRISPRCasdb data.
The "CRISPRCasdb dataset" mentioned here and in the main manuscript is composed of all groups with at least one reconstructed deletion. This dataset contains 333 groups with 4565 arrays.

#### B. Simulation details.

**Stationary length distributions** We use the stationary length distribution to sample the length of the root array for any simulation. We present the stationary length distributions we use for the independent and block deletion model.
The independent deletion length model is a $M/M/\infty$ queuing model. Spacers arrive with rate $\theta$, are served and exit after an exponential waiting time with rate $\rho_{\mathrm{I}}$. It is a well-known fact that the number of busy servers (spacers) is a Poisson distribution with rate $\frac{\theta}{\rho_{\mathrm{I}}}$, i.e.

$$p_{\mathrm{I}}(n|\theta,\rho_{\mathrm{I}}) = \left(\frac{\theta}{\rho_{\mathrm{I}}}\right)^n \frac{e^{-\frac{\theta}{\rho_{\mathrm{I}}}}}{n!}, \tag{1}$$

where $n$ is the array length. Recall, that the expected value (and variance) of a Poisson distribution is the rate, i.e. $\theta/\rho$.
The stationary length distribution of the block deletion model is more complicated to compute since there exists a dependence between different spacer positions since deletions are extended over multiple spacers. Therefore, we simplify the stationary length distribution by using the deletion rate per spacer $\rho_{\mathrm{B}} \cdot \alpha$ as parameter in a Poisson distribution and ignoring some of the complexity introduced through dependencies between spacer positions. Then the stationary length distribution of the block deletion model is a Poisson distribution with rate $\frac{\theta}{\rho_{\mathrm{B}}\alpha}$.

**Simulating the boundary effect of the block deletion model** There are two types of boundary conditions to consider for the block deletion model. The boundary effect is illustrated Figure 7 in the main manuscript.
If we assume there is a boundary effect, because repeats at the array boundary can align with less repeats, then, within the model, positions at the left or right end of the array are hit by less deletion events from the right or left, respectively. Thus, the deletion distribution across spacer positions is concave and has a fall-of near the edges of the array. To simulate this we exclude events that extend over the array border (on the left), i.e. when the sampled event length is higher than the distance to the left boundary. Due to our one-directional deletions, the effect occurs naturally on the right boundary.
If assume that there is no boundary effect, we get a uniform deletion distribution across spacer positions. In this case, since our deletions extend to the left, the last spacer is only hit by events starting there. This can be corrected by adjusting the last spacers' deletion rate to be higher. In simulations, deletions only extend to the left of the array to simplify the model, thus we need to correct the deletion rate for the rightmost spacer, since there are no events originating from the right of the spacer.

**Simulation process** First, let us recall a well-known result that helps to simplify the simulation. Let $X_1, \dots, X_n$ be independent exponentially distributed random variables with rate parameters $\lambda_1, \dots, \lambda_n$. Then

$$\min\{X_1, \dots, X_n\} \quad \textbf{(2)}$$

is also exponentially distributed, with parameter

$$\rho = \rho_1 + \dots + \rho_n. \quad \textbf{(3)}$$

Moreover, the index of the variable which achieves the minimum is distributed according to

$$P(X_k = \min\{X_1, \dots, X_n\}) = \frac{\rho_k}{\rho_1 + \dots + \rho_n}. \quad \textbf{(4)}$$

The simulation process is quite similar to the simulation process by Kupczok et al. [2].
As a first step, we simulate a coalescent tree $\mathcal{T}$ with root $r$ and choose the evolution model and its respective parameters. Then the simulation proceeds at $r$ the following way: sample from the model's stationary length distribution and generate as many spacers, i.e. create an array of natural numbers (starting from 1) with the sampled length. Set this array as the spacer array at the root $r$. We proceed by illustrating the simulation process for branches $b \in \mathcal{T}$ with respective length $t$. We continue traversing the tree in preorder, i.e. from the root towards the leafs. Since we simulated an array at $r$, each branch $b$ has an ancestor with a given spacer array $s$ of some length $n$.
Set the current time $t_c = 0$.

1. Determine the time until the next event of each type:
   (i) Draw a waiting time of the next insertion $t_\mathrm{I}$ from an exponential distribution with rate $\theta$.
   (ii) Draw a waiting time of the next deletion event $t_\mathrm{D}$ with a distribution depending on the employed deletion model. To reduce computations we rely on the result described at the start of the section. Draw the waiting time from an exponential distribution with parameter:
      (a) Independent deletion model: $\rho = n\rho_\mathrm{I}$.
      (b) Block deletion model:
         A. $\rho = n\rho_\mathrm{B}$ (simulation with boundary effect)
         B. $\rho = (n-1)\rho_\mathrm{B} + \alpha\rho_\mathrm{B}$ (simulation without boundary effect)
2. Set $t_c = t_c + \min\{t_\mathrm{I}, t_\mathrm{D}\}$. If $t_c > t$, the simulation along $b$ is finished and we return $s$ as spacer sequence of the descendant node. Else:
   (i) If $t_\mathrm{I} < t_\mathrm{D}$: Generate and insert a new spacer at the leader end of the array.
   (ii) If $t_\mathrm{D} < t_\mathrm{I}$: Determine the deletion index and delete spacers according to the selected deletion model:
      (a) Independent deletion model: Select deletion index $i$ by sampling from a uniform distribution with length $n$. Delete position $i$ from the array $s$.
      (b) Block deletion model: Select deletion index $i$ by sampling with probabilities according to Eq. (4). Then sample the length of the deletion $l$ from a geometric distribution with expected value $\alpha$. Delete the $l$ spacers starting from position $i$ to the left (or right). The boundary effect is introduced by disregarding deletions that extend beyond the first spacer in the array.
3. Continue at step 1 with the modified $s$.

**Simulation parameters** We employed the block deletion model on trees simulated according to the coalescent. Every group is composed of 16 arrays, thus every respective tree has 16 leaves. Lengths of the coalescent tree branches are randomly distributed with exponential distribution with mean $2/(k(k-1)$, where $k$ is the number of coalescing lineages.
In the independent deletion model, the following holds in the mean:

$$n \approx \frac{\theta}{\rho_{\mathrm{I}}}, \tag{5}$$

where $n$ is the mean length of the array, $\theta$ is the insertion rate and deletion rate $\rho_{\mathrm{I}}$.
Similarly, a (less precise) relationship holds for the block deletion model

$$n \approx \frac{\theta}{\rho_{\mathrm{B}}\alpha} \tag{6}$$

For all simulations for the evaluation of the performance of SpacerPlacer (Figure 5 (main manuscript) and Figures S3, S9, S8), we chose the following parameters: insertion rate $\theta = 6$, average array length $n = 22$ (the mean of the average array length of the CRISPRCasdb dataset), $\alpha \in \{1, 1.1, \dots, 4.9, 5\}$ and $\rho_{\mathrm{B}}$ was chosen such that the approximate per spacer deletion rate $\rho_{\mathrm{B}} \cdot \alpha$ remains constant, i.e. $\rho_{\mathrm{B}} = \theta/(n\alpha)$ for all $\alpha$. Then we ran 1000 simulations along 1000 sampled coalescent trees for each parameter configuration.
For the experiments, where we analyzed the contribution of unobserved spacers (Figure S8, we used the parameter estimation on the simulated data using the simulated tree and original events (no reconstruction by SpacerPlacer), excluding unobserved spacers as indicated. For the analysis of deletion distributions (Figures 7, 8 (main manuscript)) we ran 10000 simulations with the same parameters as above, but fixed $\alpha = 3$.
For the experiments shown in Figure S10, we used the (core-genome) trees and estimated parameters for the groups from the CRISPRCasdb dataset (for those with at least one deletion event). We ran 50 simulations under the BDM for each tree with the parameters estimated by SpacerPlacer for the respective group.

### Supplementary Note 2: Implementation details

**Used packages and software** SpacerPlacer is implemented in Python. We rely on Biopython [3] for the tree infrastructure during the ancestral reconstruction and tree estimation. The implementation of the joint maximum-likelihood reconstruction is based on TreeTime [4]. The visualization of the reconstruction is implemented based on ETEToolkit 3 [5] and graphs are visualized with Graphviz [6]. The multiple spacer alignment of spacer arrays is performed by MAFFT [7] using a custom script. The analytic computation of the precise likelihood function was performed with sympy [8]. The optimization of the likelihood functions for the likelihood ratio test and the parameter estimates is done with scipy [9]. For the independent deletion model we use scipy.optimize.minimize_scalar with the method "bounded" which uses Brent's Algorithm. For the block deletion model, both parameters are maximized simultaneously with scipy.optimize.minimize with the method "Nelder-Mead". We did not find that the choice of optimization method impacts the results.

**Preparation and alignment of spacer arrays** Let us consider a group of spacer arrays with at least some spacer overlap. Within each group, we identify all found unique spacers by natural numbers, i.e. to every unique spacer DNA fragment corresponds a natural number. Note, that this numbering is arbitrary. For the data of CRISPRCasdb, we performed filtering and a clustering described in Supplementary Note 1A.
We perform a Multiple Spacer Array Alignment (MSAA) of these arrays (composed of numbers!): the arrays within a group are aligned via MAFFT using the Smith-Waterson algorithm and custom match, mismatch and gap penalties. We observed that Needleman-Wunsch has similar, but slightly worse, performance. A Match is scored very highly (10), a mismatch very negatively (-10000) and the gap penalty is set to 0. Under this parameter regime, there should exist no mismatches.
For the algorithm, we convert this sequence of natural numbers into a presence-absence profile. This presence-absence profile consists of an array of 1 and 0 where each column (each element) corresponds to one spacer found in the dataset. A 1 represents that the spacer corresponding to the column is present in this individual and 0 that it is not present.
SpacerPlacer renames all multiple occurrences of the same spacer in the MSAA, i.e. their respective columns, to split them into separately evolving units within our algorithm. Thereby we guarantee the uniqueness of spacers.
This uniqueness is useful to speed up the algorithm and, since we retain the information on which spacers are identical, SpacerPlacer is able to investigate and classify the occurrence and evolution of duplicate candidates after the reconstruction step.

**Guide reconstruction** We employ the joint maximum-likelihood ancestral reconstruction algorithm introduced by Pupko et al. [10], that scales linearly with MSAA length and the number of samples, i.e. spacer arrays. Since they consider amino acids, we replaced their substitution model with a simpler model that only considers two states "spacer is present" (1) and "spacer is absent" (0) with respective insertion rate $\theta$ ($0 \to 1$) and deletion rate $\rho$ ($1 \to 0$) for the transitions. This general time reversible model corresponds to the the "unordered insertion model with independent deletions" described in the main manuscript. Following their algorithm, spacer sites are considered independent and reconstructed individually and we use their efficient method to compute the most likely states.
The implementation of the reconstruction is in large parts based on the implementation in TreeTime [4]. The insertion parameter $\theta$ and $\rho$ are chosen such that $\theta$ is much smaller than $\rho$ (default value: $\theta/\rho = 10^{-4}$). In this case, the model favors single insertions and multiple deletions over multiple insertions of the same spacer across the tree. This behavior is consistent with observations, that multiple insertions of the same spacer are rather unlikely.
We assume that the progenitor, i.e. the root, follows the same evolutionary model as the rest of the tree, instead of using a stationary length distribution. The branch length of the root is chosen as maximum over all branch lengths of the (direct) children, then the standard evolutionary model is applied to compute likelihoods. This is done to guarantee that acquisition events are sensibly placed at the root. It is possible to choose the stationary length distribution of the IDM, a Poisson distribution with parameter $\theta/\rho$, but we found it to be prone to place insertion events at the root introducing additional deletions, that could be explained more reasonably by placing insertion events on the subtrees for particular ratios of $\theta$ and $\rho$.

**Refinement process** It is useful to have a good understanding of the joint ancestral reconstruction algorithm by Pupko et al. [10] to follow the details of the refinement process.
For convenience, let us consider a single spacer position at some node $x$ with ancestral state $s_x \in \{0, 1\}$ (absence/presence) with parent node $y$ and ancestral state $s_y \in \{0, 1\}$. After constructing the guide ancestral states, we have at node $x$ the maximum likelihoods $L_x$ for the subtree $t_x$ given the state of the parent node $y$.
If a contradiction is found at $x$, we do the following: we set the most likely state $s_y$ at $y$ to 1 (presence), disregarding the computed most likely state in the guide reconstruction. This enforces that the spacer must be present at $y$. Note, that the insertion event is not guaranteed to be at $y$, only that at $y$ the spacer must be present, the insertion event can be further towards the root. Then the likelihoods are recomputed using the fixed state $s_y = 1$ which can result in differences for state $s_x$ and the whole subtree of $y$.
In this way, an insertion event is moved towards the root each time a contradiction is found. This refinement is repeated and the fixed states accumulate for each repetition until all contradictions are resolved. If the cause of the order contradiction is more than one step towards the root, a single spacer can be moved multiple times. We found that insertion events which are

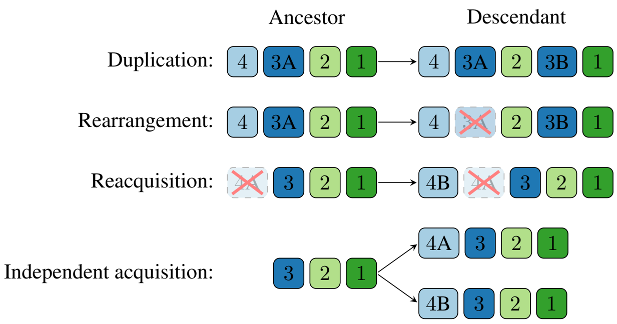


**Supplementary Fig. S1. Duplicate related event types.** We show examples for each duplicate event type that we classify. Each duplicate candidate is classified based on the evolutionary events pertaining itself and the other candidates. We classify a candidate as a *duplication*, if an insertion occurred while another candidate already exists in the array. An *rearrangement* occurs, if at the same branch the other candidate is deleted. A *reacquisition* requires that a candidate spacer was deleted earlier in time, and a new acquisition occurs at the leader end of the array. *Independent acquisitions* are, as the name implies, just independent acquisitions of the spacer (at the leader end) and no candidate existed in the array before. Independent acquisitions can for non-candidate spacers, but are generally unlikely. Any events which do not fit in any of these classes are labeled as *other duplicate related insertions*.

placed very low originally, but are very old according to the PSIO, often move multiple steps towards the root without manual placement by the above procedure, but because recomputing likelihoods returns vastly different likelihoods and ancestral states.

**Classification of spacer duplication candidates** After we obtain the MSAA, every duplicated spacer $x$ is contained in $I \in \mathbb{N}$ columns. Every such columns is individually renamed giving a set $\{x_i\}_{i=1}^{I}$ of uniquely named spacers. Since the spacer was likely acquired as a singleton at some point, there are $I-1$ duplication candidates of spacer $x$, that are likely the result of some form of duplicate related event and one "original" acquisition $x_o$.
In the following, we use a single duplicated spacer $x$ with renamed set $\{x_i\}_{i=1}^{I}$ as representative for all spacers with duplicates to describe the adjustments to the algorithm to allow the classification of the candidates.
The guide reconstruction places acquisition events for each renamed spacer $x_i$ along the tree. We adjust the subsequent reconstruction steps in the algorithm as follows:

1. Before the refinement, traverse the tree in postorder to identify the "oldest" event among the acquisitions of the $x_i$. This spacer $x_o$ can be considered the "original" spacer. It is the most confidently placed renamed spacer, because, most likely, it has the highest abundancy in the MSAA, and through its age, the most opportunities to produce contradictions to the PSIO. All $x_i$, except the "original" $x_o$ are removed from the PSIO. Through their exclusion from the PSIO potential duplicates of spacers are not subject to the contradiction treatment in the refinement step.

2. Perform PSIO-based refinement normally (without considering the candidates $x_i$ with $i \neq o$). Note that the $x_o$ are still part of the PSIO and the contradictions in which they are involved are resolved.

3. Classify the remaining candidates for duplication $x_i$ ($i \neq o$) according to their evolutionary path in the guide reconstruction. The event types are illustrated in Supplementary Fig. S1.

   Let $x_i \in \{x_i \,|\, i \neq o\}$ be the considered spacer copy and $x_j \in \{x_k \,|\, k \neq i\}$ are the other renamed spacer copies (including $x_o$) we classify $x_i$ as:

   - *Rearrangement:* A deletion event of some $x_j$ and the gain event of candidate $x_i$ are on the same branch.
     This indicates that $x$ was not duplicated but translocated. Note that inversion events (e.g. 2, 1 $\rightarrow$ 1, 2) are classified as rearrangements by this rule.
   - *Reacquisition:* A deletion event of some $x_j$ occurred not on the same branch but on the path in the tree from the root to the acquisition of $x_i$. We additionally require the acquisition of $x_i$ to be at the leader end of the array, and no other $x_j$ can be present at that time.
     The intuition is as follows: Some ancestor acquired $x$, deleted it at some point in history, and acquired the same spacer again at a later time.
   - *Duplication:* $x_i$ is inserted at a branch where some $x_j$ is already present. If the insertion of $x_i$ is at the leader end, it is instead labeled as a reacquisition.
     Here, $x$ was likely not translocated, but copied to another position in the array.
   - *Independent acquisitions:* There exist insertion events of $x_j$ in different subtrees that are not ancestral to the branch where $x_i$ was acquired.
     In this case, $x$ was likely acquired by two different samples independently.

- *Other duplicate related insertions:* If a candidate $x_i$ does not fall into any class described above, it is placed in this class.

  Ectopic insertion events, which are not rearrangements or duplications, are likely to belong to this class. Manual inspection by the user is recommended to determine the nature of the event.

**Run times of SpacerPlacer** All reconstructions, including visualizations and likelihood ratio tests, of the CRISPRCasdb dataset (518 groups) took 515.13 seconds (s) (mean per group: 0.995s, maximum: 62.76s). Of this time, 315.75s (mean per group: 0.948s, maximum: 62.76s) are used to reconstruct the 333 groups which contain at least one deletion. The forward and reverse reconstructions together (2x518 reconstructions), to determine the orientation of the arrays, took a total of 936.86s (mean per group: 0.90s, maximum: 62.76s).
Note, that these reconstructions use already provided core-genome trees. In general, we observed that tree estimation takes, by far, the most time of the reconstruction, followed by the visualization which can take a significant time for very large MSAA. As described in the data extraction section (Supplementary Note 1A), we excluded extremely large groups (4 with above 1000 spacer arrays) since their run time is substantial (their runtime can exceed the complete run time of the rest of the dataset) and they have limited impact on parameter estimates.

### Supplementary Note 3: Spacer array based tree estimation

Although independently estimated phylogenies should be preferred for the reconstruction of ancestral CRISPR arrays, SpacerPlacer also offers an alternative approach. It allows for the estimation of trees solely based on the spacer arrays themselves in cases where there is no independent estimate of the phylogeny available or when the phylogeny of the CRISPR loci is anticipated to deviate from the core phylogeny.
In this case, SpacerPlacer can estimate a tree using an Unweighted Pair Group Method with Arithmetic mean (UPGMA) or Neighbor Joining (NJ) approach based on a custom spacer based distance for each pair of spacer arrays.
This distance function is a simple likelihood based distance function that employs a straightforward evolutionary model in between two (related) arrays and their ancestor. It follows a block deletion model with ordered insertions characterized by three parameters the insertion rate $\theta$, the deletion initiation rate $\rho_{\mathrm{B}}$ and mean deletion length $\alpha$.
The distance between two arrays $a_1, a_2$ with common ancestor $a$, with distances $t_1, t_2$ from $a_1$ and $a_2$ respectively, is computed as follows:

1. Find the first equal spacer (FES) of arrays $a_1, a_2$.

2. We split spacers into younger and older spacers than the FES, i.e. in front/behind the FES. The motivation is, that the FES is the youngest spacer found in $a$, excluding e.g. the possibility of ectopic insertions or unobserved spacers.

   Then we consider younger spacers (in front of the FES) as insertions on the branch from ancestor to $a_1$ or $a_2$ respectively, while older spacers (behind the FES) are assumed to be present at the ancestor and deleted or retained along the branches according to their absence or presence in $a_1$ and $a_2$.

   To compute the likelihoods of the insertions and deletion events determined above, we use the following evolutionary model

   - For insertions, we use a poisson distribution with mean $\frac{\theta}{\rho\alpha}$ dependent on branch lengths $t_1, t_2$ (see Supplementary Note 1B). This is only a rough approximation of the insertion mechanism.
   - For deletions we use the block deletion model with parameters $\rho_{\mathrm{B}}$, $\alpha$, dependent on branch lengths $t_1, t_2$. The likelihoods are computed according to the simplified or precise likelihood function computed in Supplementary Note 4.

3. We maximize this likelihood function with respect to $t_1, t_2$, treating $\theta, \rho_{\mathrm{B}}, \alpha$ as constants, resulting in $t_{1,\max}, t_{2,\max}$. The distance between $a_1$ and $a_2$ is then given by $t_{1,\max} + t_{2,\max}$. Maximum and minimum branch lengths are bounded for numerical stability and to retain minimal branch lengths to allow event placement for the maximum likelihood reconstruction.

Then, the tree is constructed with UPGMA with respect to this distance function and the pairwise distances between the given set of arrays.
SpacerPlacer provides suitable parameters $\theta, \rho_{\mathrm{B}}$ and $\alpha$ estimated on CRISPRCasdb data for the spacer based tree reconstruction. Custom parameters can be chosen by the user.
We do not suggest to use spacer based trees to estimate model parameters. If ancestral reconstructions are estimated based on these trees, the estimates of parameters will be scaled according to the scale of the parameters ($\theta$, $\rho_{\mathrm{B}}$, $\alpha$) used for the tree reconstruction.

### Supplementary Note 4: Derivation of the likelihood functions, bias-corrections and the likelihood ratio test

To compute parameter estimates and compare our models with a likelihood ratio test we compute the likelihood functions of both the independent deletion model (IDM) and the block deletion model (BDM). Note, that the independent deletion model is a special case of the block deletion model ($\alpha = 1$). Nevertheless, we find it instructive to discuss the IDM separately to illustrate the differences between both models.

To compute the likelihood functions, the main issue to solve is that a connected component of consecutive spacers that have been lost along a branch of the phylogeny can arise from a combination of multiple block deletion events of different lengths. SpacerPlacer provides two ways to compute the combined likelihood for each of these connected components. On the one hand, we provide a simplified approach (see Section B) that approximates the likelihood, ignoring the exact time and sequence of possible combinations of different block deletion events along a branch. On the other hand, in a more precise approach (see Section C), we take into account the order and timing of the events. This gives rise to several ordinary differential equations that can be solved analytically, but solving them is computationally expensive.

Let us define the notation: We denote the underlying tree as $\mathcal{T}$, the tree $\mathcal{T}$ is composed of a finite collection of branches $b_i, i \in I$ for some set $I \subset \mathbb{N}$. We denote the length of a branch $t_b$. We denote the set of lengths of adjacent deletions (gaps) of branch $b$ as $K_b := \{k_i\}_{i=1}^{m_b}$, where each $k_i$ denotes the length of adjacent deletions that are separated by at least one retained spacer and $m_b \in \mathbb{N}$ is the total number of these sets along branch $b$. We write $|K_b| := \sum_{i=1}^{m_b} k_i$ for the total number of deleted spacers along branch $b$. The collection of all deletion sets along $\mathcal{T}$ is denoted by $\mathcal{K} := \{K_b \,|\, b \in \mathcal{T}\}$ Furthermore, we denote for each branch $b$ the number of spacers present at the parent node $N_b$ and the set of all these numbers $\mathcal{N} := \{N_b\}_{b \in \mathcal{T}}$ along $\mathcal{T}$.

**A. Independent deletion model.** In the independent deletion model (IDM) the likelihood of deletions is independent from their position in the array, and in particular, their adjacency.

The model is governed by one parameter, the deletion rate $\rho_\mathrm{I}$ at each site in the alignment. The likelihood along each branch $b$ is solely determined by $\rho_\mathrm{I}$, the branch length $t_b$, the number of deletions $|K_b|$ and the number of retained spacers, i.e. spacers present in the parent and not deleted in the child, which is given by $N_b - |K_b|$. Since the waiting times for deletions at each site are independent and exponentially distributed the probability of retaining a spacer $P(1 \to 1)$ is given by

$$P(1 \to 1) = e^{-\rho t_b}, \tag{7}$$

and the probability of deleting a spacer $P(1 \to 0)$ by

$$P(1 \to 0) = 1 - P(1 \to 1) = 1 - e^{-\rho_\mathrm{I} t_b} \tag{8}$$

for each branch $b$, where $t_b$ denotes the branch length. Since all spacer sites are independent the likelihood of losing $|K_b|$ spacers and retaining $N_b - |K_b|$ spacers along branch $b$ with deletion rate $\rho_\mathrm{I}$ is given by

$$L_\mathrm{I}(b, N_b, K_b, \rho_\mathrm{I}) = e^{-\rho_\mathrm{I} t_b (N_b - |K_b|)} \left(1 - e^{-\rho_\mathrm{I} t_b}\right)^{|K_b|}. \tag{9}$$

Since the events on different branches are independent, the likelihood of the collection of deletions $\mathcal{K}$ occurring along $\mathcal{T}$ is

$$\mathcal{L}_\mathrm{I}(\mathcal{T}, \mathcal{N}, \mathcal{K}, \rho_\mathrm{I}) := \prod_{b \in \mathcal{T}} L_\mathrm{I}(b, N_b, K_b, \rho_\mathrm{I}). \tag{10}$$

**B. Simplified likelihood function of the block deletion model.** The computation of the likelihood function of the block deletion model (BDM) is non-trivial. We compute a simplified version of this likelihood function and, a more complex, more precise likelihood function (see Section C). We rely mainly on the simplified version of the likelihood function of the BDM, due to its stronger performance in parameter estimations of simulations (compare Fig. S3 and Fig. S9) and relative ease of computation. For more information about the simplification and differences to the more precise computation see section C.

The BDM is governed by two parameters, the deletion initiation rate $\rho_\mathrm{B}$ and a parameter $\alpha$ determining the average deletion length. Each deletion event of any length still has a probability of $P(1 \to 0) = 1 - e^{-\rho_\mathrm{B} t_b}$ to occur. We assume the length of deletions to be geometrically distributed, i.e. $g_\alpha(k) := \mathrm{Geom}(\alpha^{-1}) = (1 - \alpha^{-1})^{k-1} \alpha^{-1}$ is the probability of a block deletion to have the length $k$.

We observe that deletions separated by a retained spacer are independent. Thus we can consider each set of adjacent deletions separately. For each adjacent deletion of some length $k$, we have to consider all combinations of events that sum up to a total deletion length $k$. As an example consider a deletion of length 3 under the block deletion model. It can either be three independently occurring single deletion events of length 1, one deletion event of length 3, one of length 2 and one of length 1, or one of length 1 and one of length 2.

By some further consideration of the general case to delete a joint set of length $k \geq 1$, we obtain a recursive formula for the probability. The probability of deleting a joint set of length $k$ in blocks is given by the recursive function

$$f(k,\rho_{\mathrm{B}},\alpha) = (1-e^{-\rho_{\mathrm{B}} t_b}) \left( \sum_{j=1}^{k} f(k-j,\rho_{\mathrm{B}},\alpha) g_{\alpha}(j) \right), \tag{11}$$

$$f(0,\rho_{\mathrm{B}},\alpha) = 1. \tag{12}$$

The intuition behind $f$ is that we divide each deletion $k$ into all possible subsets of deletions of lengths $1,\dots k$, adding up to $k$ in total, and then consider these subsets as independent deletions of respective lengths $k_i$. Then, these subsets are divided into possible subsets of length $1,\dots k_i$, and so on, until no subdivision is possible (i.e. only a length 1 block remains). Of course, each single deletion event needs to be initiated, which produces the term $P(1 \to 0) = (1-e^{-\rho_{\mathrm{B}} t_b})$.
It is illustrative to compute the first components of $f$, for notational brevity we omit the parameters $\rho_{\mathrm{B}}, \alpha$, then

$$\begin{aligned}
f(0) &= 1,\\
f(1) &= P(1\to 0)g(1),\\
f(2) &= P(1\to 0)\left(f(1)g(1)+g(2)\right)\\
&= P(1\to 0)^2 g(1)^2 + P(1\to 0)g(2)\\
f(3) &= P(1\to 0)\left(f(2)g(1)+f(1)g(2)+g(3)\right)\\
&= P(1\to 0)^3 g(1)^3 + 2P(1\to 0)^2 g(1)g(2) + P(1\to 0)g(3)\\
f(4) &= P(1\to 0)\left(f(3)g(1)+f(2)g(2)+f(1)g(3)+g(4)\right)\\
&= P(1\to 0)^4 g(1)^4 + 3P(1\to 0)^3 g(1)^2 g(2) + 2P(1\to 0)^2 g(1)g(3) + P(1\to 0)^2 g(2)^2 + P(1\to 0)g(4).
\end{aligned}$$

In short, the computation of $f(k)$ is an exercise of combinatorics of all possibilities to produce a deletion of length $k$. The powers of $g(j)$ and $P(1\to 0)$ are the number of required events of length $j$ to produce the whole deletion of length $k$ (in this specific configuration). The multiplicative constants are produced by the different possible compositions of $k$ (such as a deletion of length 3 being composed of deletion events of length $\{1,2\}$, $\{2,1\}$).
For the likelihood of a branch $b$ with adjacent deletion $K_b := \{k_i\}_{i=1}^{m_b}$ we need to include the probabilities for retained spacers, as in the IDM case. Thus we have for the likelihood of branch $b$

$$L_{\mathrm{B}}(b,N_b,K_b,\rho_{\mathrm{B}},\alpha) = e^{-\rho_{\mathrm{B}} t_b (N_b - |K_b|)} \prod_{i=1}^{m_b} f(k_i,\rho_{\mathrm{B}},\alpha). \tag{13}$$

Since the events on different branches are independent, the likelihood of $\mathcal{T}$ is given by

$$\mathcal{L}_{\mathrm{B}}(\mathcal{T},\mathcal{N},\mathcal{K},\rho_{\mathrm{B}},\alpha) = \prod_{b\in\mathcal{T}} L_{\mathrm{B}}(b,N_b,K_b,\rho_{\mathrm{B}},\alpha). \tag{14}$$

**C. Precise likelihood of the block deletion model.** As the name implies, the simplified block deletion model makes some strong assumptions to ease computations. While treating deletions separated by surviving spacers and branches as independent is correct, the function $f$, i.e. the probability of losing adjacent losses of length $k$, is much more difficult to compute in general.
In the *simplified* block deletion model we consider the events that compose a deletion as independent, each deletion event happens with rate $\rho_{\mathrm{B}}$ and contains at least one spacer.
The simplified likelihood function only considers the sequencing of events. As an example see Figure S2 (a), there we only consider that the deletion of length 5 can be composed of events of different length and can be differently sequenced, e.g. $\{3,2\},\{2,3\},\{1,1,3\},\{1,3,1\},\dots$.
We disregard two important facts that are illustrated in Supplementary Fig. S2. Firstly, we disregard the *event times*. For accurate computation of the likelihood we have to keep in mind that it is a continuous process with random jumps where the sequencing of events *and* their event times are significant.
Secondly, we dismiss that *no other* deletion events are allowed to happen on the retained spacers until the deletion event occurs. Then, if a deletion occurs at time $b_e$, only the remaining spacers need to be considered for the likelihood computations along the remaining branch length $b - b_e$, since already deleted spacers do not contribute to the likelihood anymore.
For example, in Fig. S2, if a deletion of length 3 occurs first at time $j_1$, 4 spacers need to be *not* hit by a deletion event during the same time and, after $j_1$, one spacer needs not to be hit till the second event occurs at time $j_2$. If events are reversed, then the likelihoods till the first event $j_1$ remains the same, *but* after the deletion 2 spacers need to be retained till $j_2$ hits. Thus, there is much more complexity in computing possible configurations and their likelihoods compared to the simplified approach. Keep

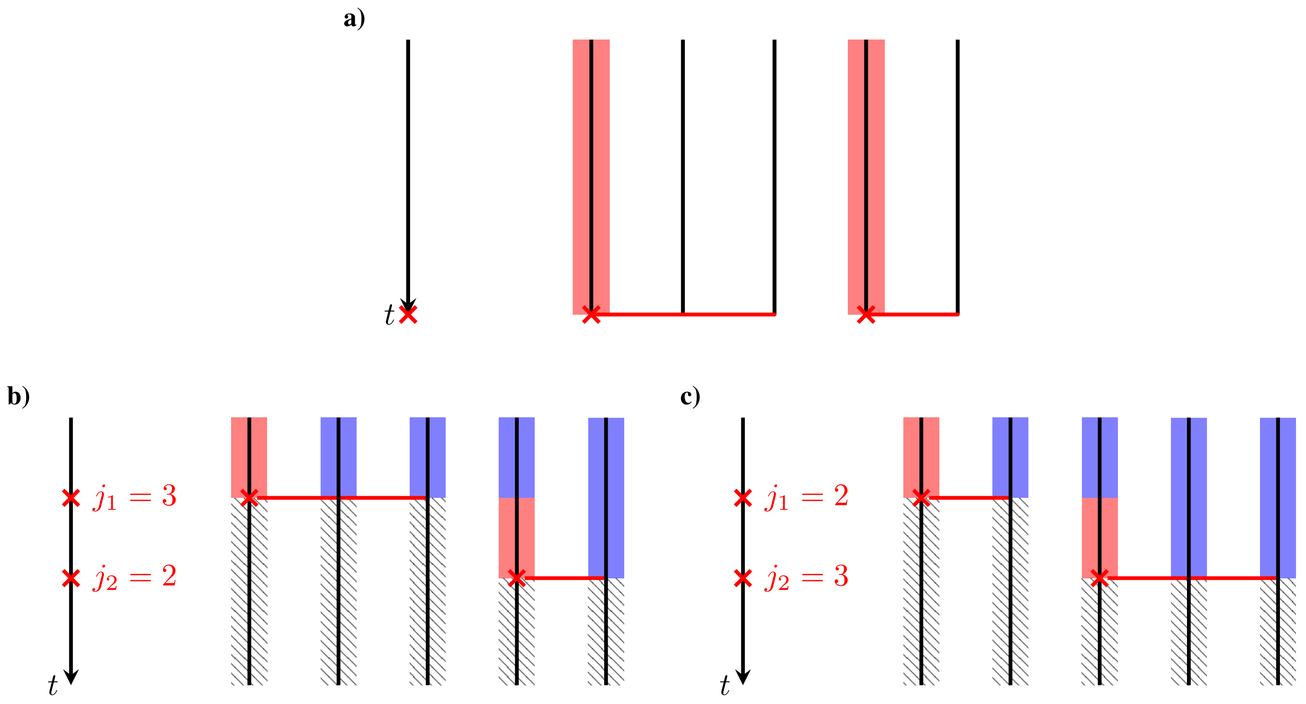


**Supplementary Fig. S2. a)** illustrates the naive approach employed for computing the simplified likelihood of the block deletion model. Each line corresponds to one spacer position that is deleted in time $t$, i.e. we consider a deletion of length 5. Red shading marks the time in which a deletion event can happen. Crosses indicate the start of a deletion event, and the red line indicates how many positions are hit. Note that in **a)** we do not consider any positions beside the marked two positions. The figure shows only one realization of a deletion of length 5. For the complete likelihood all possible configurations of deletion lengths that add up to 5, and their sequencing, need to be considered.
**b)** and **c)** show two possible realizations of a deletion of length 5 and illustrates how one needs to think about deletions to compute the precise likelihood function. Red shading still marks the time in which a deletion event might hit a specific spacer position. Different from **a)** we need to consider how long other spacer positions have to be preserved, which is indicated by blue shading. Hatched area indicates time where the spacer is deleted, i.e. they do not contribute to the likelihood anymore. Be aware that the event times $j_1, j_2$ are exponentially distributed, and thus require a continuous time approach to compute the likelihood. Furthermore, the sequencing of events is relevant due to a differing number of positions that need to be preserved (i.e. the blue area is different).

in mind that the event times are exponentially distributed as well. Fortunately, very similar problems have been investigated previously [11]. In our case, we are confronted with a special case of a *continuous time Markov chain* known as *pure death process*.

A *pure death process* is a continuous time, discrete state, Markov process. More precisely, we have a family of random variables $\{X(t) \,|\, 0 \leq t < \infty\}$ dependent on time $t$ where the possible values of $X(t)$ are positive integers, e.g. a population of individuals, in our case the length of a deletion block. Since we consider a Markov process, we have a transition probability function

$$P_{ij}(t) = P(X(t+u) = j | X(u) = i), \quad i,j = 0,1,2,\dots$$

that is independent of $u$. This function describes the probability of getting from $i$ individuals to $j$ individuals in time $t$ starting from time $u$. Since we consider a pure death process, we assume that $P_{ij} = 0$ for all $j > i$, i.e. no individuals are birthed and individuals can only die.

By making some careful considerations, one can come up with a ordinary differential equation describing $P_{ij}$ [11].

Often, the pure death process is considered with the restriction that in each infinitesimal time-step only one individual can die, i.e. $P_{ij} \neq 0$ for $i,j$ satisfying $|i-j| = 1$, and $P_{ij} = 0$ for $i,j$ with $|i-j| > 1$.

In our case, we consider a spacer block of length $k$ and want to know the probability that at time $t$ no spacer remains, i.e. the transitional probability $P_{k0}(t)$. But since we allow the deletion of multiple spacers described by a geometric distribution, we need to consider a more involved differential equation.

By adapting the considerations in [11], we construct the initial value problem

$$P'_{k0}(t) = \rho_{\mathrm{B}} \sum_{j=0}^{i-1} g_\alpha(i-j)(j+1)\left(P_{j0}(t) - P_{i0}(t)\right), \tag{15}$$

$$P_{00}(t) = 1, \tag{16}$$

where $P'_{ij}$ is the derivative of $P_{ij}$ with respect to the time $t$, $g_\alpha$ is the geometric distribution and $\rho_{\mathrm{B}}, \alpha$ are the block deletion model parameters.

We are only interested in the probability to delete all spacers $P_{k0}(t)$ in time $t$ and thus do not state the equation in full generality for $P_{ij}$.

Let us explore this equation a bit further. We delete spacer blocks with constant rate $\rho_{\mathrm{B}}$, in that sense we remain close to the (standard) pure death process, but the length of our losses is geometrically distributed leading to a more involved equation, since it is possible to jump from any value between $k$ and 1 to 0.

This implies, as can be seen in Eq. (15), that $P'_{k0}$ depends on all $P_{j0}$ where $j < k$. This means, to compute $P_{k0}$ for some $k$ requires us to solve all previous ordinary differential equations for $j < k$. Clearly, the initial condition $P_{00} = 1$ arises naturally for our model, as an empty set of spacers can not lose additional spacers.

Solving these equations analytically (and numerically) is possible, but, through the sequential dependency, gets more difficult for increasing $k$ quickly. We computed the likelihood function analytically up to a maximum joint deletion length of $k_{\text{max}} = 68$. Since CRISPR arrays are generally short in length this suffices for most of the groups in our datasets.

In case of a larger set of adjacent deletions of size $k > k_{\text{max}}$, we split the deletion into $z$ fragments with length $k_{\text{max}}$ and a residual of length $k_{\text{res}}$ such that $k = z \cdot k_{\text{max}} + k_{\text{res}}$. We regard these $(z+1)$ fragments as independent and compute their likelihood accordingly.

Remarkably, as can be seen in Supplementary Fig. S3 compared to Supplementary Fig. S9 the parameter estimates computed with the simplified function (and no bias corrections) yield better estimates then the ones computed with the precise likelihood function.

Moreover, the computation is more expensive than the simplified version. We provide the option to use this likelihood function in SpacerPlacer, but we recommend to use the simplified likelihood function (with bias corrections) for most applications.

### D. Correcting biases of the likelihood functions.

**Correcting the unobserved bias** In the main manuscript, we discussed that the estimates fall short with respect to unobserved spacers, i.e. events that occur somewhere within the tree, but are not observed in the data since the spacer was deleted along the whole tree. We consider the IDM case to illustrate this correction. At the end of the section we give a short description how we handle the bias correction for the BDM.

To compute the corrected likelihood, we normalize the likelihood of observing the events characterized by $N_b, K_b, \rho_{\mathrm{I}}$ at branch $b$ by the likelihood that all acquired spacers (at branch $b$) are observable, i.e. all acquired spacers were *not* deleted along the subtree of $b$.

Thus, we first compute the likelihood $\xi_{\mathrm{I}}^{\text{unobs}}(b, \rho_{\mathrm{I}})$ that one spacer present before branch $b$, i.e. inserted at the parent branch, is unobservable along the subtree, recursively. Let $C_b$ be the children of $b$, then

$$\xi_{\mathrm{I}}^{\text{unobs}}(b,\rho_{\mathrm{I}}) = P(1 \to 0, b, \rho_{\mathrm{I}}) + P(1 \to 1, b, \rho_{\mathrm{I}}) \prod_{b_c \in C_b} \xi_{\mathrm{I}}^{\text{unobs}}(b_c, \rho_{\mathrm{I}}), \tag{17}$$

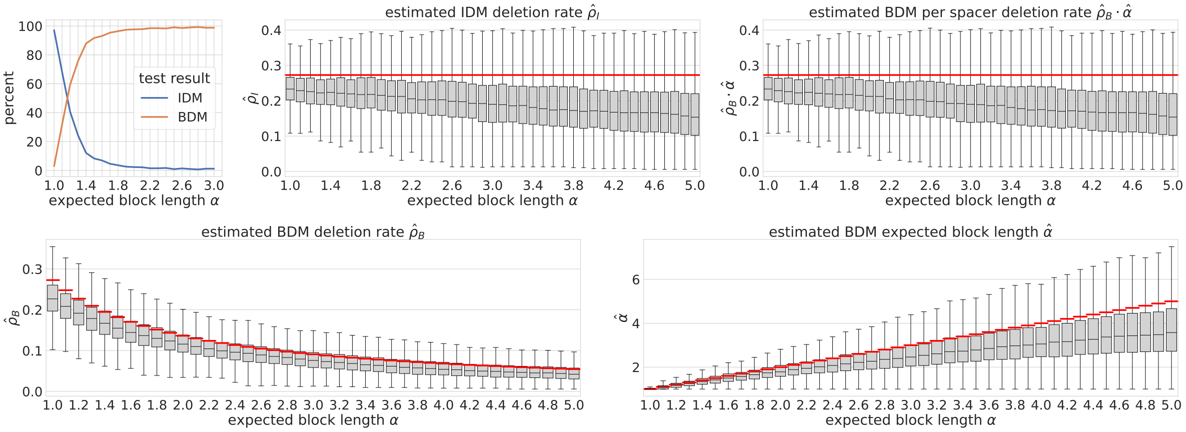


**Supplementary Fig. S3.** Likelihood ratio test performance and parameter estimates of simulations using the precise likelihood function. The true parameters used for the simulations are shown as red lines. The decreased performance compared to the simplified likelihood function Supplementary Figure S9) suggests that either the precise likelihood function is not as numerically stable, or inherent biases are even stronger compared to the simplified likelihood function.

where at terminal branches $b_{\text{ter}}$ we have

$$\xi_{\text{I}}^{\text{unobs}}(b_{\text{ter}}, \rho_{\text{I}}) = P(1 \to 0, b_{\text{ter}}, \rho_{\text{I}}). \tag{18}$$

Since an insertion at branch $b$ needs to be deleted along all child branches (and can't be deleted at the insertion branch), the likelihood $L_{\text{I}}^{\text{unobs}}(b, \rho_{\text{I}})$ for a single spacer inserted at branch $b$ to remain unobserved is given by

$$L_{\text{I}}^{\text{unobs}}(b, \rho_{\text{I}}) := \prod_{b_c \in C_b} \xi_I^{\text{unobs}}(b_c, \rho_{\text{I}}). \tag{19}$$

Then the corrected likelihood, conditioned on observing the $g_b$ spacers acquired at each branch $b$ in the sample, is given by

$$\frac{L_{\text{I}}(b, N_b, K_b, \rho_{\text{I}})}{\left(1 - L_{\text{I}}^{\text{unobs}}(b_c, \rho_{\text{I}})\right)^{g_b}}, \tag{20}$$

since we assume the spacer positions to be independent.
And finally we obtain the corrected likelihood for the whole tree

$$\frac{\mathcal{L}_{\text{I}}(\mathcal{T}, \mathcal{N}, \mathcal{K}, \rho_{\text{I}})}{\prod_{b \in \mathcal{T}} \left(1 - L_{\text{I}}^{\text{unobs}}(b, \rho_{\text{I}})\right)^{g_b}}. \tag{21}$$

By optimizing this likelihood, we obtain a bias corrected maximum likelihood estimate of the deletion rate $\rho_{\text{I}}^{\text{corr}}$ of the IDM. Since in the BDM, deletions started at unobserved spacers introduce a dependency that is very hard to quantify (and computationally expensive), it is infeasible to adjust the BDM estimate precisely. Thus, we approximate the bias correction of $\hat{\rho}_{\text{B}}^{\text{corr}}$ by using the bias correction of the IDM. In formulas, we compute $\Delta\hat{\rho}_{\text{I}} = \hat{\rho}_{\text{I}}^{\text{corr}} - \hat{\rho}_{\text{I}}^{\text{no corr}}$ to obtain a corrected estimate $\hat{\rho}_{\text{B}}^{\text{corr}} = \hat{\rho}_{\text{B}} + \Delta\hat{\rho}_{\text{I}}$ of the BDM deletion rate.

**Correcting the bias of the geometric distribution** The maximum likelihood estimator of the geometric distribution is known to be biased (especially for a low number of samples). Let $n := |\mathcal{K}|$ be the number of deletions along the tree $\mathcal{T}$ and $\hat{p} = 1/\hat{\alpha}$, then we correct the estimation by computing

$$\hat{p}^{\text{corr}} = -\frac{1}{2}\left(\sqrt{(n+1)^2 - 4n\hat{p}} - n + 2\hat{p} - 1\right), \tag{22}$$

and we obtain the bias corrected estimator $\hat{\alpha}^{\text{corr}} = 1/(\hat{p} + \hat{p}^{\text{corr}})$.

### E. Maximum likelihood ratio test.

Here, we give a detailed description of the likelihood ratio test introduced in the main manuscript to assess the goodness of fit of the IDM and BDM. Note, that the IDM and BDM are nested, since the IDM is a special case ($\alpha = 1$) of the BDM. We define our hypotheses

$$H_0: \quad (\rho, \alpha) \in \Theta_0, \tag{23}$$

$$H_1: \quad (\rho, \alpha) \in \Theta \setminus \Theta_0, \tag{24}$$

where $\Theta_0 := [0,\infty) \times \{1\}$ and $\Theta := [0,\infty) \times [1,\infty)$, and we can write down the likelihood ratio test statistic

$$\lambda_{\text{LR}} := \frac{\sup\limits_{\rho_{\text{I}} \in [0,\infty)} \mathcal{L}_{\text{I}}(\mathcal{T},\mathcal{N},\mathcal{K},\rho_{\text{I}})}{\sup\limits_{(\rho_{\text{B}},\alpha) \in \Theta} \mathcal{L}_{\text{B}}(\mathcal{T},\mathcal{N},\mathcal{K},\rho_{\text{B}},\alpha)} = \frac{\sup\limits_{\rho_{\text{I}} \in [0,\infty)} \prod_{b \in \mathcal{T}} L_{\text{I}}(b,N_b,K_b,\rho_{\text{I}})}{\sup\limits_{(\rho_{\text{B}},\alpha) \in \Theta} \prod_{b \in \mathcal{T}} L_{\text{B}}(b,N_b,K_b,\rho_{\text{B}},\alpha)} = \frac{\sup\limits_{\rho_{\text{I}} \in [0,\infty)} \prod_{b \in \mathcal{T}} L_{\text{B}}(b,N_b,K_b,\rho_{\text{I}},1)}{\sup\limits_{(\rho_{\text{B}},\alpha) \in \Theta} \prod_{b \in \mathcal{T}} L_{\text{B}}(b,N_b,K_b,\rho_{\text{B}},\alpha)}, \quad \textbf{(25)}$$

The maximization can be simplified and computed more efficiently by taking the logarithm. Furthermore, by Wilks' theorem

$$-2\ln(\lambda_{\text{LR}}) \sim \chi^2(1), \quad \textbf{(26)}$$

asymptotically, where $\chi^2(n)$ is the Chi-squared distribution for $n$ degrees of freedom (in our case $n = 1$, since $\alpha$ is the only degree of freedom). Note, that if there is no deletion at all ($\mathcal{K} = \emptyset$), we have $L_{\text{I}}(b,N_b,0,\rho_{\text{I}}) = e^{-\rho_{\text{I}} b N_b}$, $f(\rho_{\text{B}},\alpha,0) = 1$, $L_{\text{B}}(b,N_b,0,\rho_{\text{B}},\alpha) = e^{-\rho_{\text{B}} b N_b}$ for all $b \in \mathcal{T}$ and thus $\lambda_{\text{LR}} = 1$ with maximum points $\hat{\rho}_{\text{I}} = \hat{\rho}_{\text{B}} = 0$ ($\hat{\alpha} \in [1,\infty)$).
Note, that we do not use bias corrections for the likelihood ratio test (only for the parameter estimates).

| Task | SpacerPlacer | Kupczok et al. [2] | Collins et al. [19] |
|---|---|---|---|
| Phylogeny computation | external or<br>blockwise UPGMA/NJ | rooted NJ | external or<br>most parsimonious |
| Guide Reconstruction | unordered acq.<br>individual del. | — | custom acq. costs<br>custom del. costs |
| Refined Reconstruction | partially ordered acq.<br>individual del. | | |
| Parameter estimation | ordered acq.*<br>individual del.<br>blockwise del. | ordered acq.<br>individual del.<br>fragment del. | — |
| Deletion models for LRT | individual vs blockwise | — | — |

**Supplementary Table S1.** The table shows a comparison of the spacer deletion and insertion models used here and in related publications [2, 19]. Naturally, the most parsimony approach by Collins and Whitaker [19] does not include rates for different events but custom cost functions for ordered and ectopic acquisitions, duplications, deletions depending on their position and length.
*The parameter estimation and the likelihood ratio testing in SpacerPlacer rely solely on reconstructed deletions, thus the acquisition rate can only be estimated indirectly.

### Supplementary Note 5: Comparison to other models of spacer array evolution

Some stochastic models have been specifically designed for CRISPR spacer array evolution [2, 12], while others explored the coevolutionary dynamics between phages and bacteria [13–15]. Moreover, the length of spacer arrays has been the subject of multiple modeling approaches [16, 17]. The equilibrium between spacer deletions and acquisitions will lead to an average spacer array length, which could be subject to selection [16]. In our setting with no fitness effect of spacers, the number of spacers in an array is Poisson distributed and depends on the ratio of overall acquisition and per spacer deletion rate.

Kupczok and Bollback compared models of spacer deletions that assume independent deletion of single spacers and polarized insertions with models with internal insertions, as well as a model in which fragments of the spacer array could be deleted [2]. In contrast to the block deletions with geometrically distributed length considered here, fragment deletions cover, regardless of its length, any possible fragment of the array with the same probability. Based on the data available at the time, Kupczok and Bollback found that independent deletions, as well as fragment deletions are plausible models. The block deletion model proposed here has two advantages: a) the observed deletion pattern observed in CRISPRCasdb and experiments [18] are in line with block deletions of a geometrically distributed length and b) the memorylessness of the geometric distribution allows efficient likelihood computation for reconstruction and parameter estimation. In the past, the independent deletion model has been used to construct a deletion rate estimator that is based only on the variability at the trailer-end of spacer arrays [12], but did not incorporate the other parts of the array nor ancestral reconstructions.

Collins and Whitaker proposed an alternative to parametric models by introducing a reconstruction tool based on maximum parsimony principles [19]. To accommodate all possible types of events in CRISPR array evolution, they defined empirically determined scores for acquisitions at the leader end, duplications, insertions within the array, deletions (of arbitrary length), and independent acquisitions.

In contrast, we rely on explicit evolutionary models that can offer insight beyond ancestral reconstruction through parameters that correspond to molecular mechanisms and evolutionary processes. We use a dual approach where the underlying molecular mechanisms of CRISPR arrays influenced the modeling choices, but the reconstruction-based parameter estimates also allows investigating the evolutionary (co-)dynamics of CRISPR arrays. For example, only our new framework allows us to introduce the likelihood ratio test that identifies the mode of deletions.

An overview of the models used in related studies and different steps of our pipeline, including our LRT workflow, is shown in Supplementary Table S1.

### Supplementary Note 6: Applications

**A. CRISPR based tree estimation in the medical setting.** Due to CRISPR arrays' fast spacer turnover compared to the mutation rate of genomes in genral, tree estimation based on the CRISPR array allows to track small timescale phylogenetic relationships of samples. SpacerPlacer allows this small timescale tracking through its ability to construct phylogenetic trees from CRISPR array diversity.

In [20], Tomida et al. proposed to study phylogenetic relationships based on CRISPR spacer array composition compared to Multi Locus Sequence Typing (MLST) to get trees of higher resolution between the closely related hospital samples.

They analyzed CRISPR arrays of *Helicobacter cinaedi* found in 6 different hospitals in Japan with some additional reference samples from other countries. They identified CRISPR arrays and identified their spacer content. Depending on CRISPR genotype they could be separated in different related groups.

Tomida et al. constructed MLST trees and found them to be unsatisfying since their resolution was not high enough to distinguish even between very distant in time samples (Supplementary Fig. S4a)).

As an alternative, they reconstructed trees based on a pairwise distance between CRISPR arrays and UPGMA which allows them to distinguish the ST-4/CC4 samples (Supplementary Fig. S4b)). But the Hospital A samples seem to be separated by the foreign and fairly old samples which suggests that the might be different from the real underlying ancestral relations between the samples. Furthermore, we do not expect the pairwise distance to be well suited to estimate CRISPR based trees, since it does not respect block deletions and the insertion order.

We show in Supplementary Fig. S4c) that a evolutionary approach, as employed by SpacerPlacer, is more suitable, since Hospital A samples are placed closely together and the foreign samples are placed as outgroups. Additionally, we are able to reconstruct the ancestral history of the CRISPR arrays with SpacerPlacer. We used CCTK [19] on the same dataset. The results are shown in Supplementary Fig. S4d). It returns a similar tree to SpacerPlacer with the main differences between the older samples from outside of Japan.

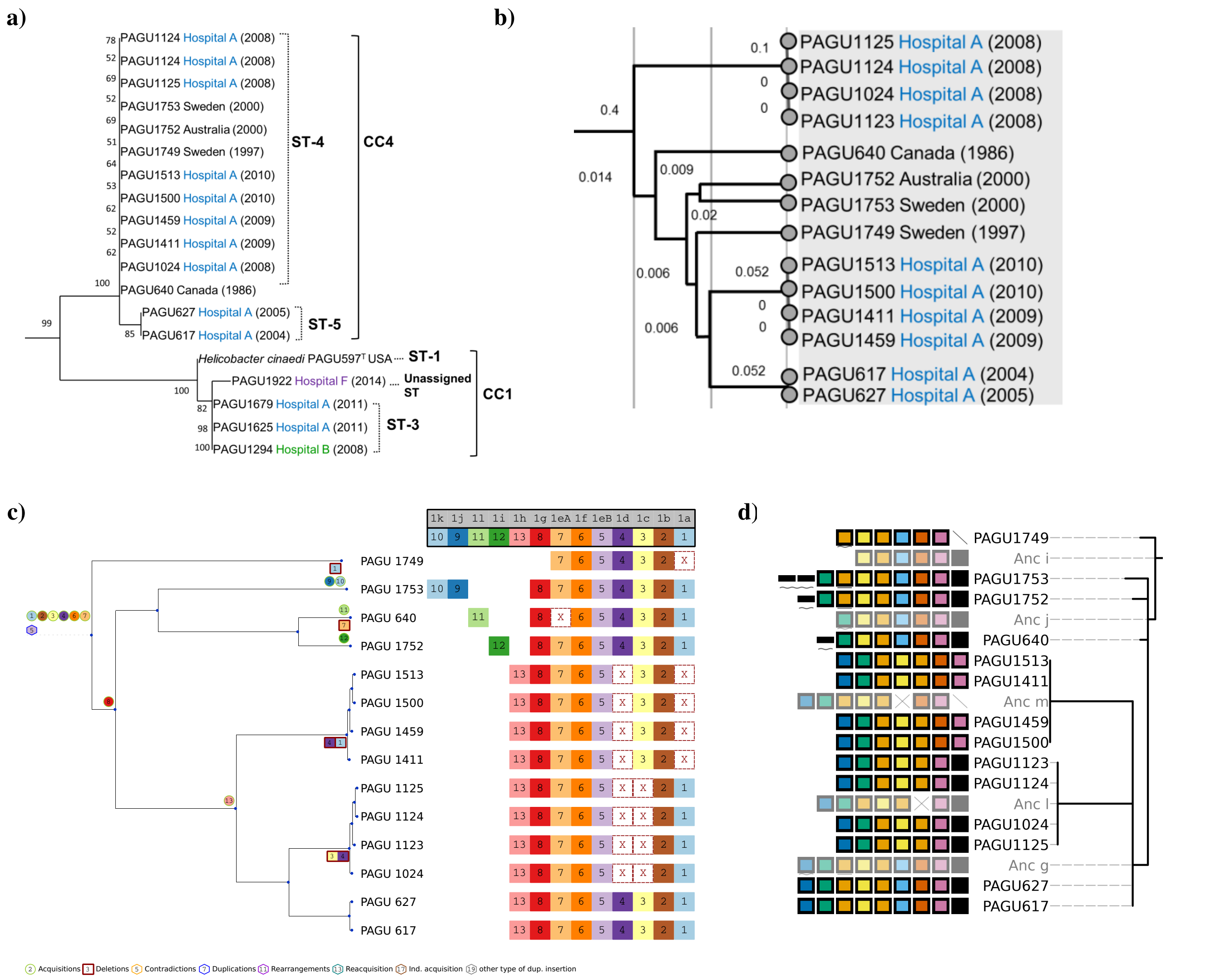


**Supplementary Fig. S4.** This is a comparison between trees produced by different methods by Tomida et al. and SpacerPlacer for the same subset of data [20].

**a)**: Shows a tree reconstructed by Tomida et al. with MLST. Note, that within the group CC4 almost no distinction can be made, only ST-5 can be separated from ST-4.

**b)**: Shows a UPGMA tree based on pairwise distance between the CRISPR arrays (on spacer level). As can be seen, the variance of the array spacer content is enough to construct a non-trivial tree, but we would expect the older samples of foreign origin (PAGU 640, PAGU1752, PAGU1753, PAGU1749) to be an outgroup of this tree.

**c)**: Shows a reconstruction of both the tree and the ancestral states with SpacerPlacer of the same group of CRISPR arrays. Note, that within this tree: all Hospital A samples are placed closely together and samples of foreign origin (PAGU 640, PAGU1752, PAGU1753, PAGU1749) are placed together as outgroup.

**d)**: Shows a reconstruction of both tree and ancestral states with CCTK [19] with default parameters. Similarly, to c) the samples of foreign origin are placed together as outgroup. Hospital A arrays that are close in time are placed closely together.

**B. CRISPR array dynamics from long-read microbiome data.** The quality of our reconstruction of CRISPR evolution depends heavily on the quality of available data. We require long contiguous sections of CRISPR arrays, preferably the whole CRISPR array, to determine the exact spacer order.

If only small snips of CRISPR arrays are available, as is often the case in microbiome data, no complete order can be determined and single CRISPR arrays might be split across multiple samples. Reconstruction of incomplete reads of CRISPR arrays with SpacerPlacer is certain to distort the estimations of trees and evolutionary parameters.

Lam and Ye characterize CRISPR arrays from a gut microbiome sequenced with long-read sequencing technologies [21]. Through the use of spacer alignments and spacer graphs they analyzed the dynamics of spacer acquisition within their datasets. Although they used data composed of long reads, many of their reads do not contain complete CRISPR arrays or arrays with uncertain ends. Furthermore, some groups show only limited diversity in the spacer arrays.

We show the dataset named "group5" by Lam and Ye. It is composed of complete reads of CRISPR arrays and shows enough diversity to produce a non-trivial ancestral reconstruction.

The reconstruction of "group5" by SpacerPlacer shown in Supplementary Fig. S5, indicates the potential of estimating a phylogenetic tree based on CRISPR arrays. On a local scale the tree reconstruction performs well, producing clusters that have close evolutionary relations. However, the estimated tree produces quite a few independent acquisitions (e.g. spacer 11, 109, 118, 120, ...), which suggests to us, that some of the samples are placed too far apart, even though they have a quite close evolutionary relations.

We believe that this issue is produced by our use of the relatively simple reconstruction method UPGMA that only operates on pairwise distances. We expect that the CRISPR array based tree could be much improved, especially on non-local scale, by more sophisticated approaches to tree reconstruction and refinement. We also used CCTK [19] to estimate a tree and produce an ancestral reconstruction. The results are similar.

a)

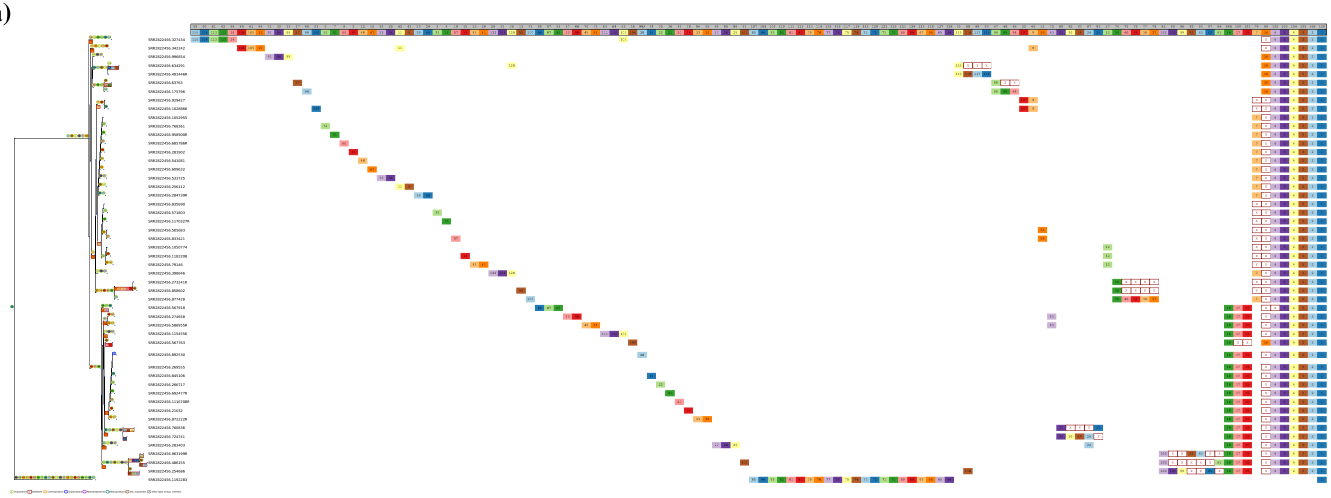

b)

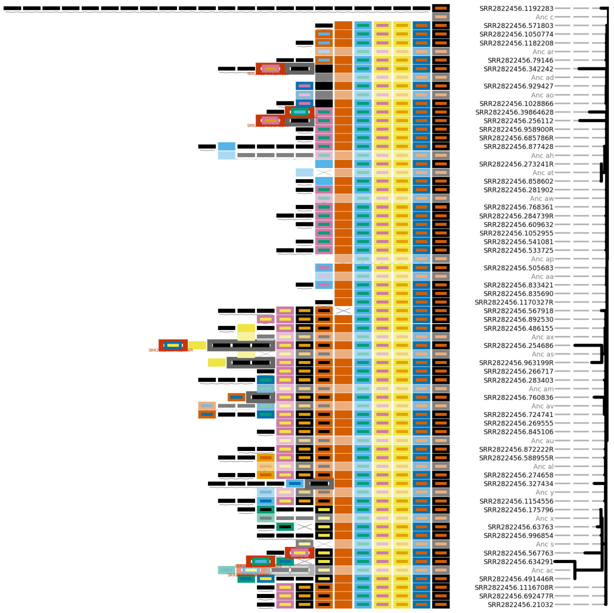


**Supplementary Fig. S5. a)**: Reconstruction of Lam and Ye's "group5" with SpacerPlacer. The tree is estimated with SpacerPlacer. **b)**: Reconstruction and tree of the same dataset estimated with CCTK [19] using default parameters.
In both figures, some arrays which contain exactly the same spacers were combined together to not unnecessarily bloat the figures.

**a)**

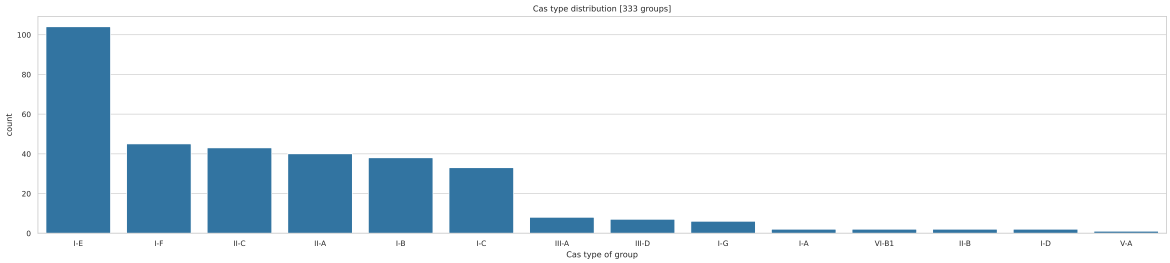


**b)**

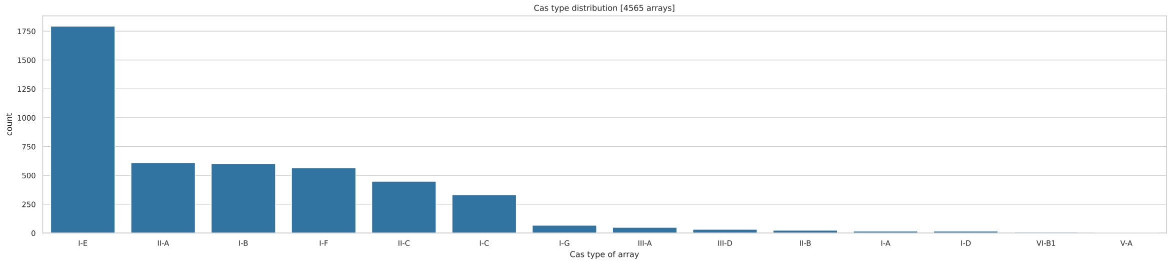


**Supplementary Fig. S6.** We show the count of groups (**a)**) and spacer arrays (**b)**) according to their Cas type of the CRISPRCasdb dataset. Note that groups are composed of arrays of the same Cas type.

**a)**

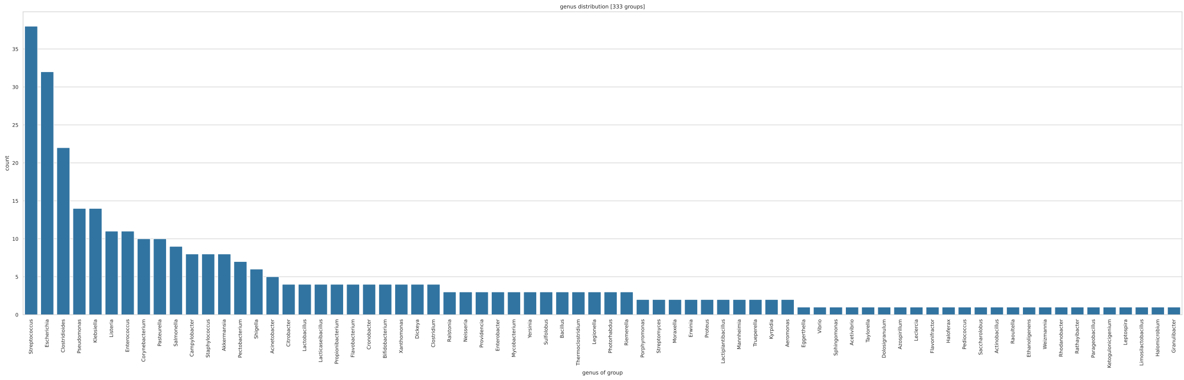


**b)**

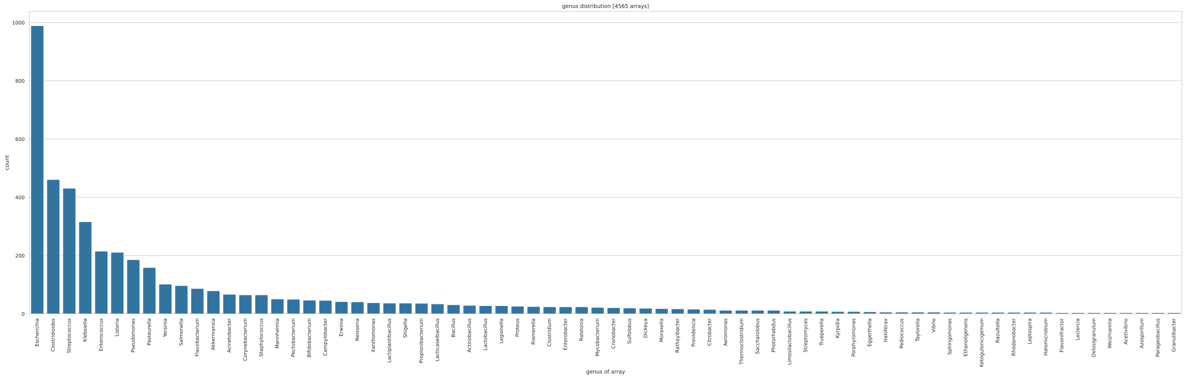


**Supplementary Fig. S7.** We show the genus counts of groups (**a)**) and of arrays (**b)**) of the CRISPRCasdb dataset. Note that groups are composed of arrays of the same genus.

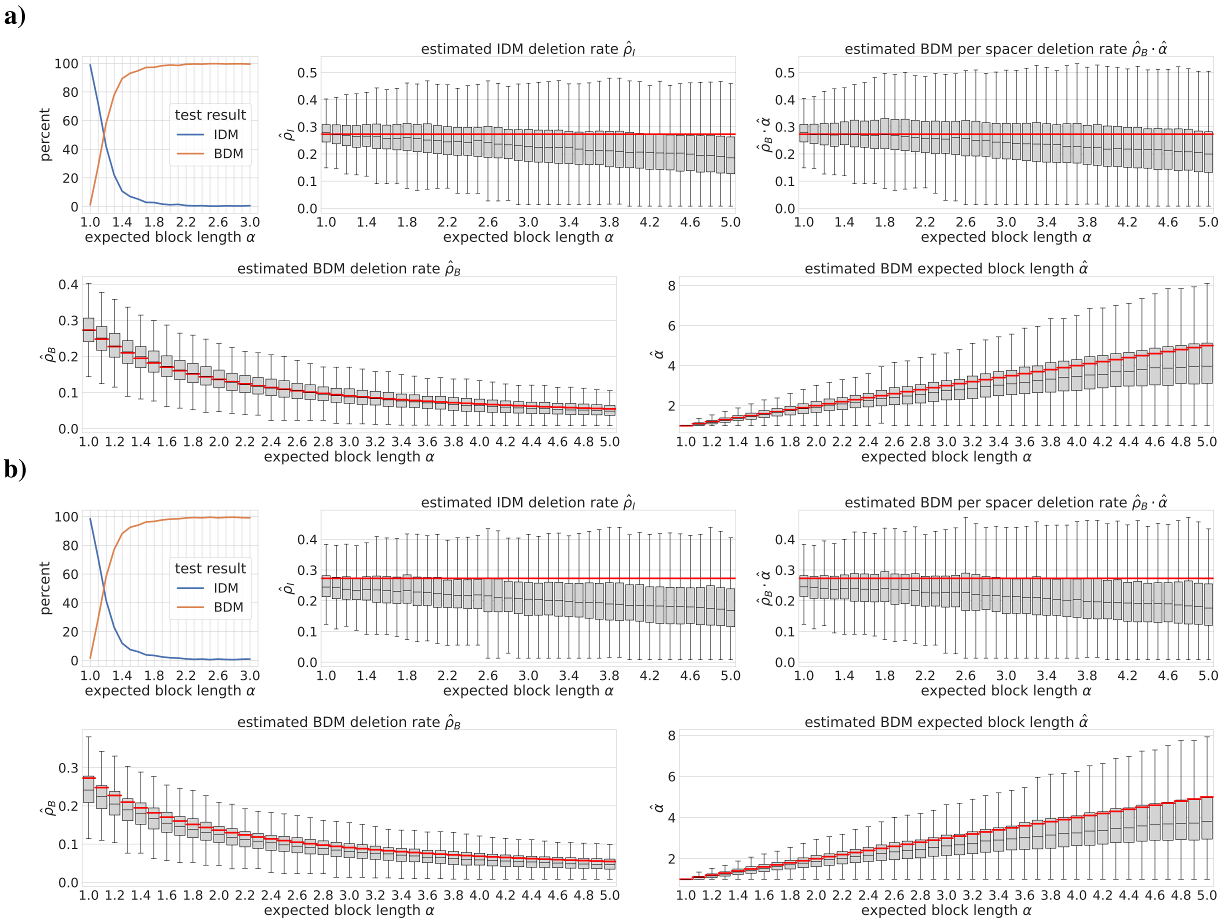


**Supplementary Fig. S8.** We show parameter estimates of simulations where we used the original simulated events as the reconstruction. In **a)** unobserved spacers are part of the reconstruction. Note, that same branch insertions and deletions of the same spacer are not included in the reconstruction, since all computations of the algorithm are branch based, while simulations work in a continuous time setting. In **b)** we rely on the original simulated events, excluding all events pertaining unobserved spacers. No bias corrections are performed which leads to an underestimation of mean deletion length $\alpha$ in particular. The decrease of performance in **b)** illustrates the large effect of unobserved spacers on parameter estimates. This effect can largely be alleviated by bias corrections (see Figure 5 (main manuscript)).

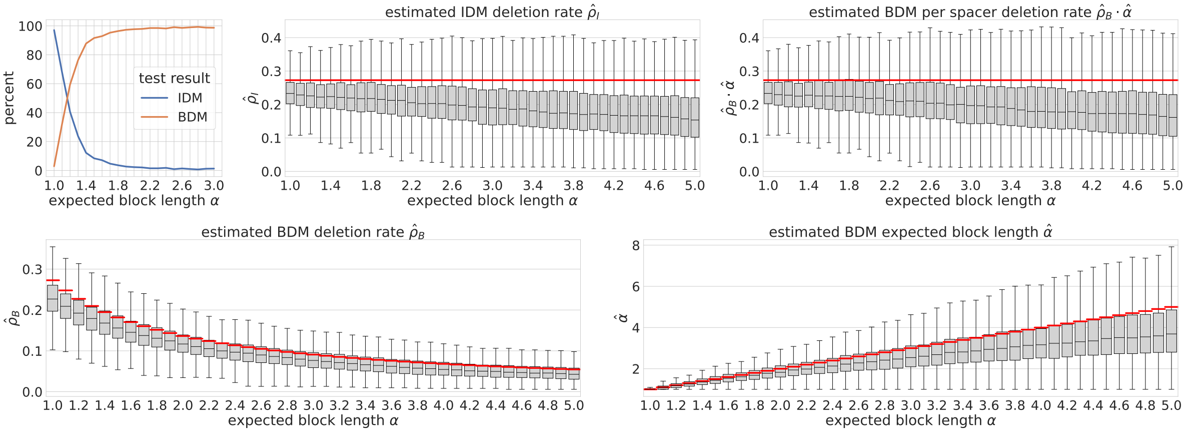


**Supplementary Fig. S9.** Likelihood ratio test performance and parameter estimates of simulations using the simplified likelihood function *without* bias corrections for the deletion rates $\rho_I, \rho_B$ and mean deletion length $\alpha$. The true parameters used for the simulations are shown as red lines. Note, the better performance compared to estimates based on the precise likelihood function shown in Figure S3.

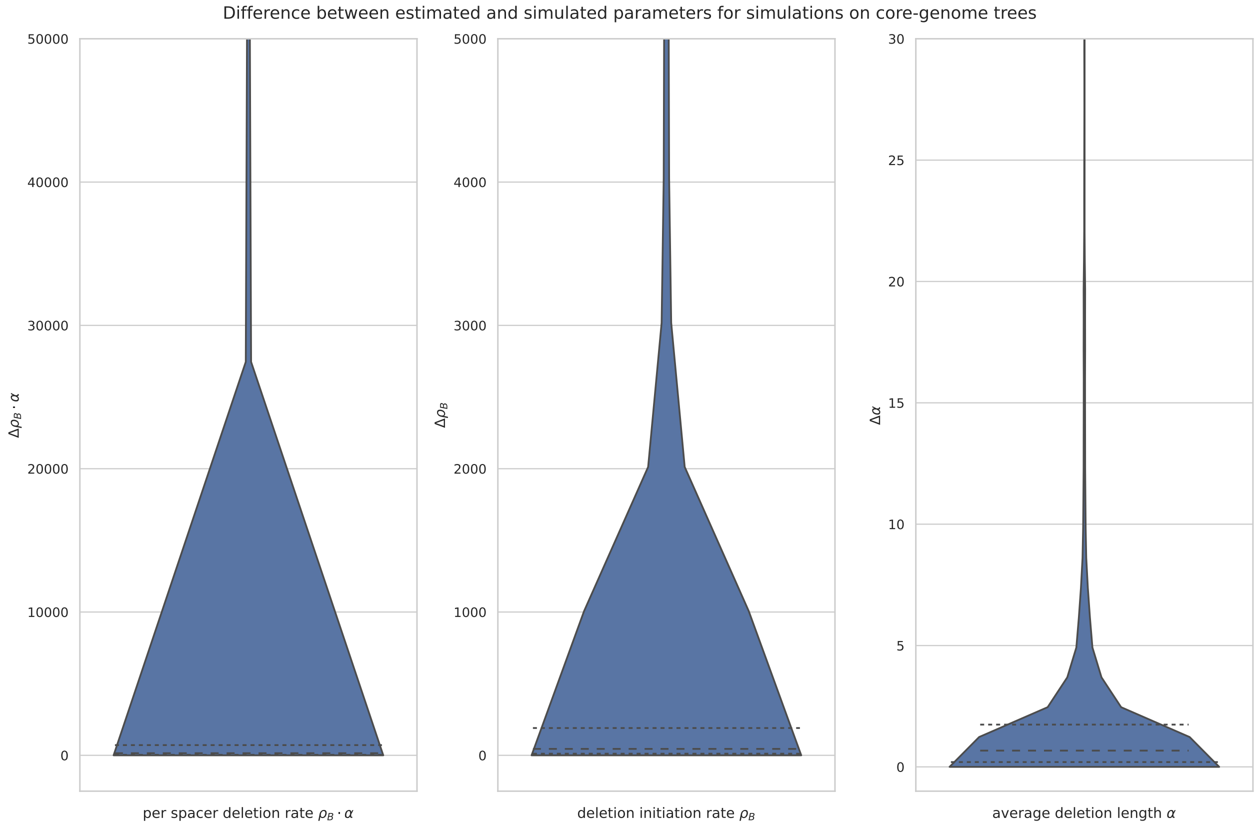


**Supplementary Fig. S10.** We show the parameter estimates on simulations on the core-gene based trees of the CRISPRCasdb dataset with parameters chosen according to the SpacerPlacer estimates. For each group we ran 50 simulations under the BDM on the respective tree using the parameter estimates by SpacerPlacer as simulation parameters.

In each plot, the (absolute) difference between the ground truth, i.e. the simulation parameters, and the estimate based on the reconstruction by SpacerPlacer. We limited the y-axis for readability. The dashed lines show the quartiles.

As can be seen, medians of the estimates are quite precise. Outliers arise mostly for datasets with few leaves where the quality of the tree, and the estimated timescale, is questionable. Furthermore, the estimates on the real data have high variance, are dependent on small samples and are not always reliable. Thus, as discussed in the main manuscript, we believe that the underlying tree is of crucial importance for good performance. Moreover, parameter estimates for single groups should be considered with caution and we suggest to use large datasets to obtain parameter estimates in the median. Note, that the mean can be heavily skewed by outliers.

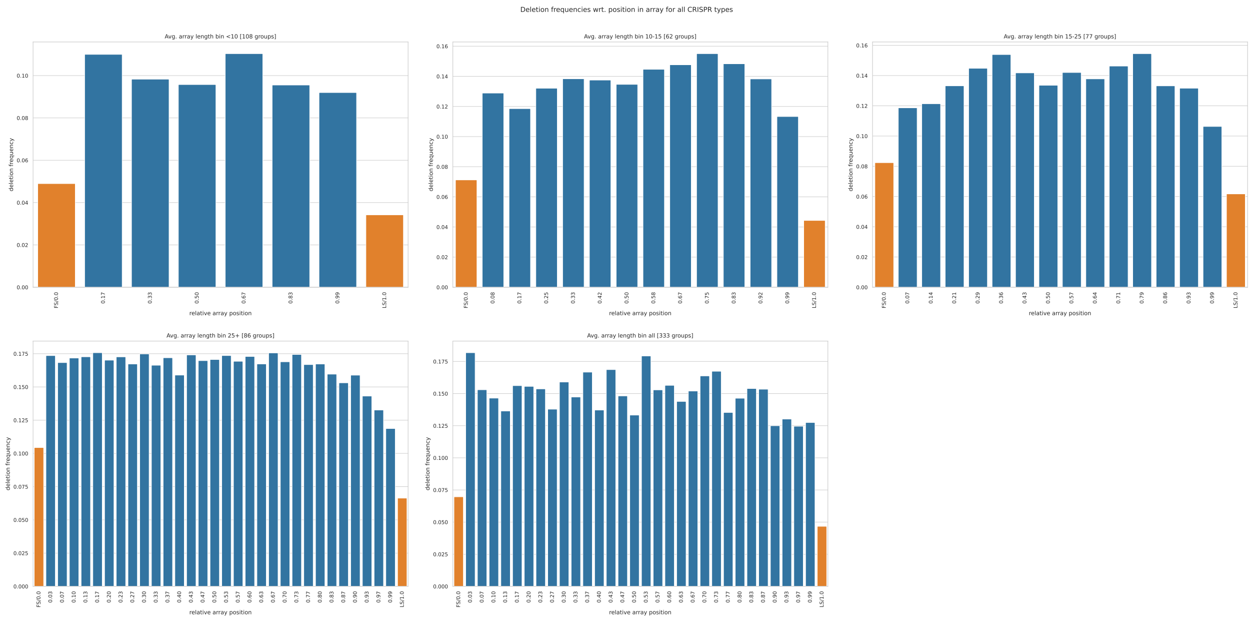


**Supplementary Fig. S11.** We show the deletion frequencies for spacers at different positions within the array for groups within different ranges of average array length (at the leafs) in the CRISPRCasdb dataset. The number of groups considered in each plot is shown in each figures title. Deletions were counted and binned according to their position relative to the array length at the visited branch. Note, that much of the variance between close bins is the result of the binning process, since arrays differ in length along the reconstructed trees. Indicated in orange are the first (0.0) and last (1.0) spacers positions as separate bins.

#### Simulated deletions along the array

#### Reconstructed deletions along the array

**1a)** array boundary does not affect deletion block length

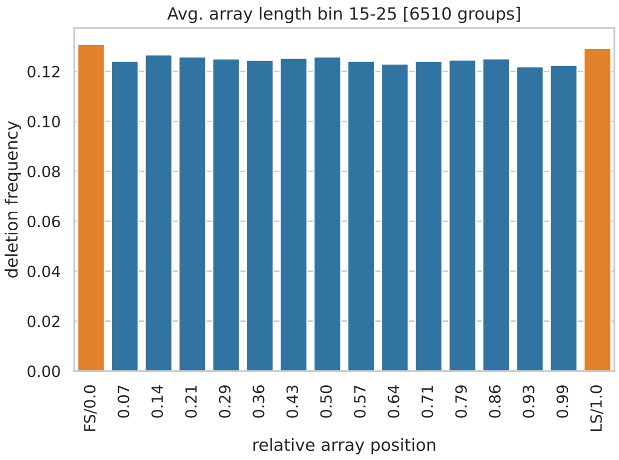


**1b)** reconstructed from simulations shown in 1a

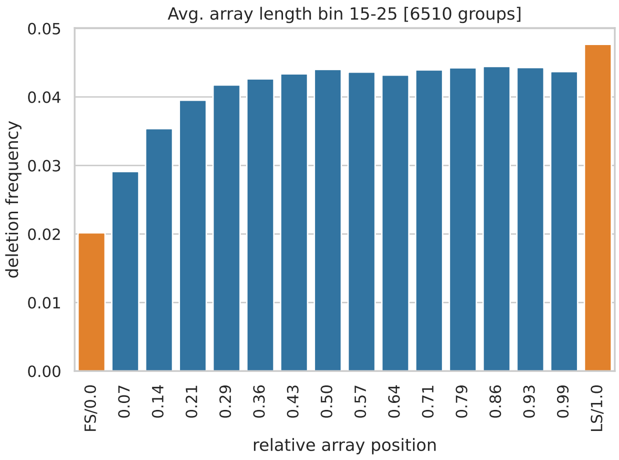


**2a)** boundary affects deletion block length

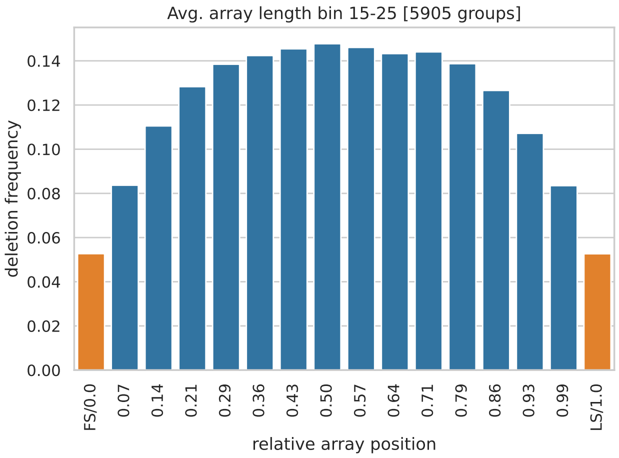


**2b)** reconstructed from simulations shown in 2a

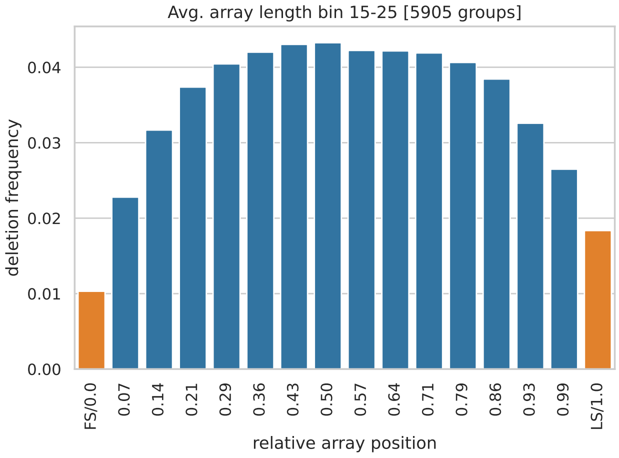


**3)** reconstructed from CRISPRCasdb data

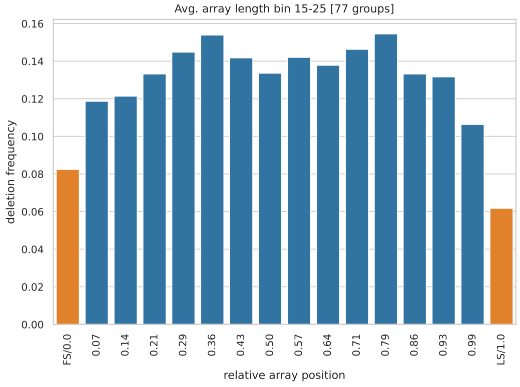


**Supplementary Fig. S12. Deletion frequency with respect to the position along the array.** The bars display the proportion of branches in the tree where a deletions occurs for different relative positions along the array as indicated by the x-axis. **1a)** shows the deletion frequencies for simulations under an artificial block deletion model, where deletions of internal spacers and spacers at the ends of the array occur at the same frequency. In contrast, **2a)** shows the deletion frequencies for simulations under the more natural block deletion model where block deletions cannot exceed the array boundary as nearby repeats need to align. Here, deletions are naturally less likely to occur towards both ends. **1b)** and **2b)** show the corresponding values based on the events reconstructed by SpacerPlacer. In both cases SpacerPlacer is able to reconstruct the deletion distribution of the respective models, except near the leader end of the array. This result is not surprising, as errors in the reconstruction of recently acquired spacers are naturally less likely to be corrected by the PSIO. In reconstructions based on CRISPRCasdb data, **3)**, the deletion frequencies show a distribution similar to the deletion model with a boundary effect (2a and 2b). But there are more reconstructed deletions at the first spacer position than at the last spacer position, contrasting the pattern we would expect based on the neutral simulations.
For each plot, we bin all deletion events according to their relative position compared to the current length of the array at the parent node of each branch. Each bar in the figures corresponds to a bin, with the label on the x-axis indicating the upper limit of the bin. The first spacer (FS/0.0) and the last spacer (LS/1.0) are placed in their own categories (shown in orange).

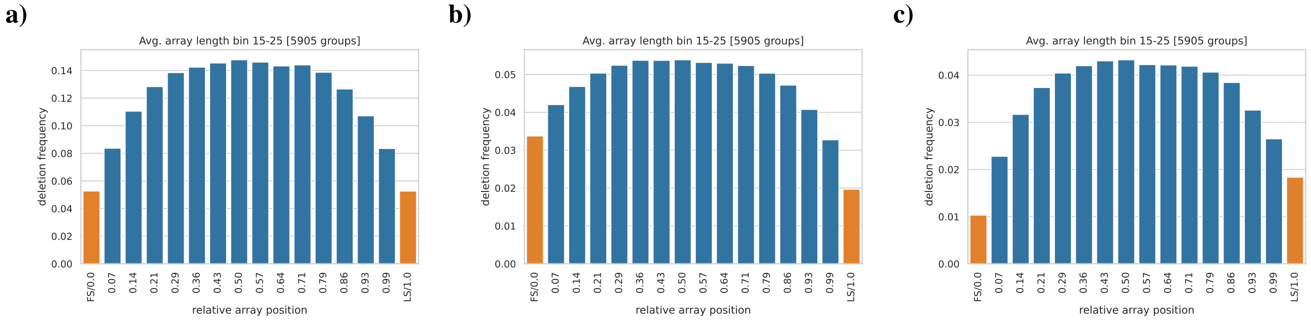


**Supplementary Fig. S13.** We compare the effect of branch based versus event based determination of deletion frequency in simulations. **a)** shows the deletion frequency distribution along the relative array position for simulations with boundary effect. This graph is based on the relative positions at each deletion event time.

**b)** shows the deletion frequency distribution, if we only use branch based information as is the case for reconstructions by SpacerPlacer. Since SpacerPlacer does not estimate event times, the relative array position of spacers is determined by their position in the parent node. The relative array position of a spacer changes during the evolution along the branch due to insertions and deletions. Thus, further along the branch, the first spacer might not be the first spacer anymore, which means that deletion events started at younger, newly inserted spacers can hit the former first spacer. This results in increased deletion rates for the first position bin and some subsequent position bins. Furthermore, **a)** and **b)** are scaled differently. **a)** is scaled by the actual number of deletion events, while **b)** is scaled by the number of nodes in the tree.

**c)** shows the reconstruction by SpacerPlacer of the simulated data. As can be seen, the deletion rates at the start of the array are strongly underestimated relative to the end of the array. This reconstruction effect dominates the deletion rate increase caused by the reduction of branch based information. **a)** and **c)** are part of Supplementary Fig. S12.

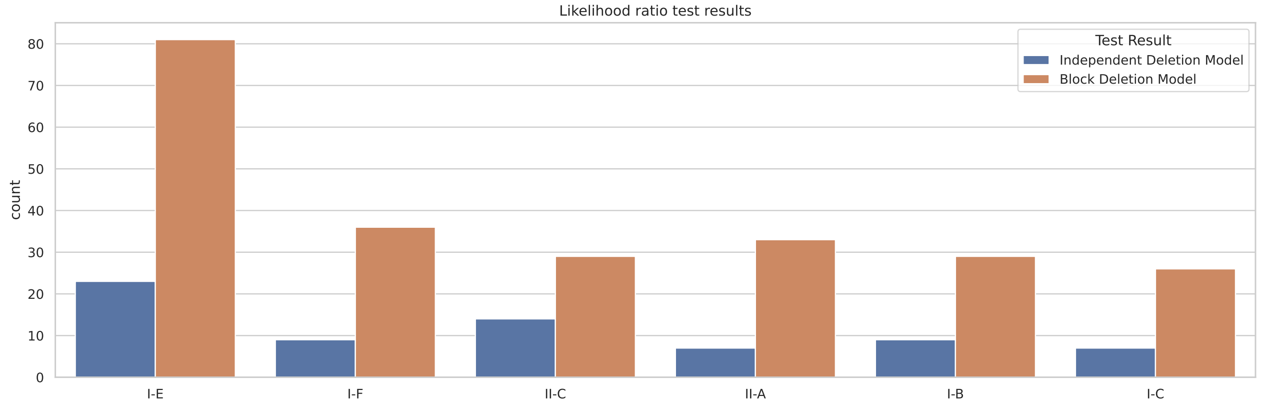


**Supplementary Fig. S14.** Likelihood ratio test results of CRISPRCasdb data. The significance level was chosen as $0.05$. As can be seen, the test decides against the null hypothesis, i.e. the independent deletion model, a significant amount of times for all Cas types. We only show Cas types were a substantial number of groups are available in the dataset.

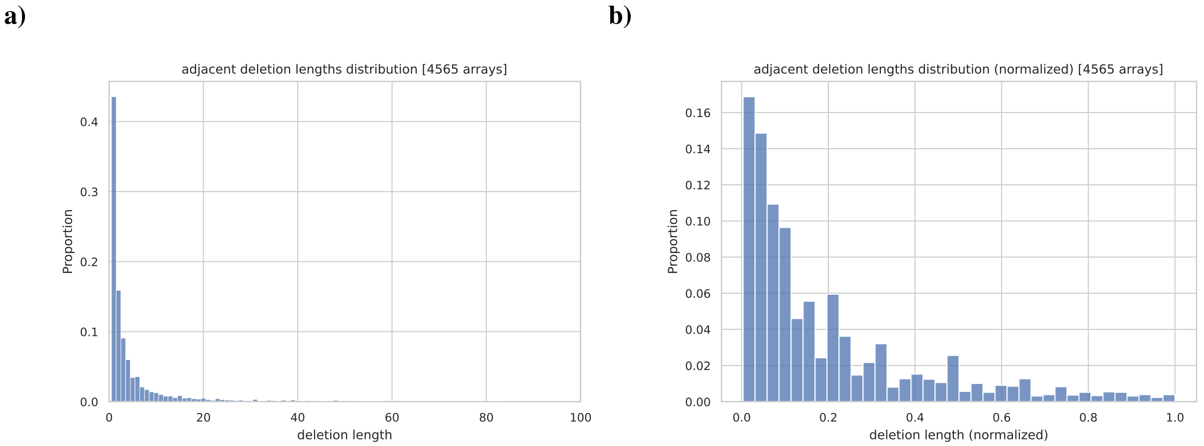


**Supplementary Fig. S15.** **a)** shows the distribution of the block lengths of adjacent deletions observed in reconstructions of the CRISPRCasdb dataset. **b)** shows the distribution normalized to the length of the parent node of the branch where the deletion occurred. The distribution roughly follows a geometric distribution, but there exist more long deletions than would be expected under a geometric distribution. Note, that the shown block deletion lengths show the "gap" lengths and could be composed of multiple deletions of shorter length.

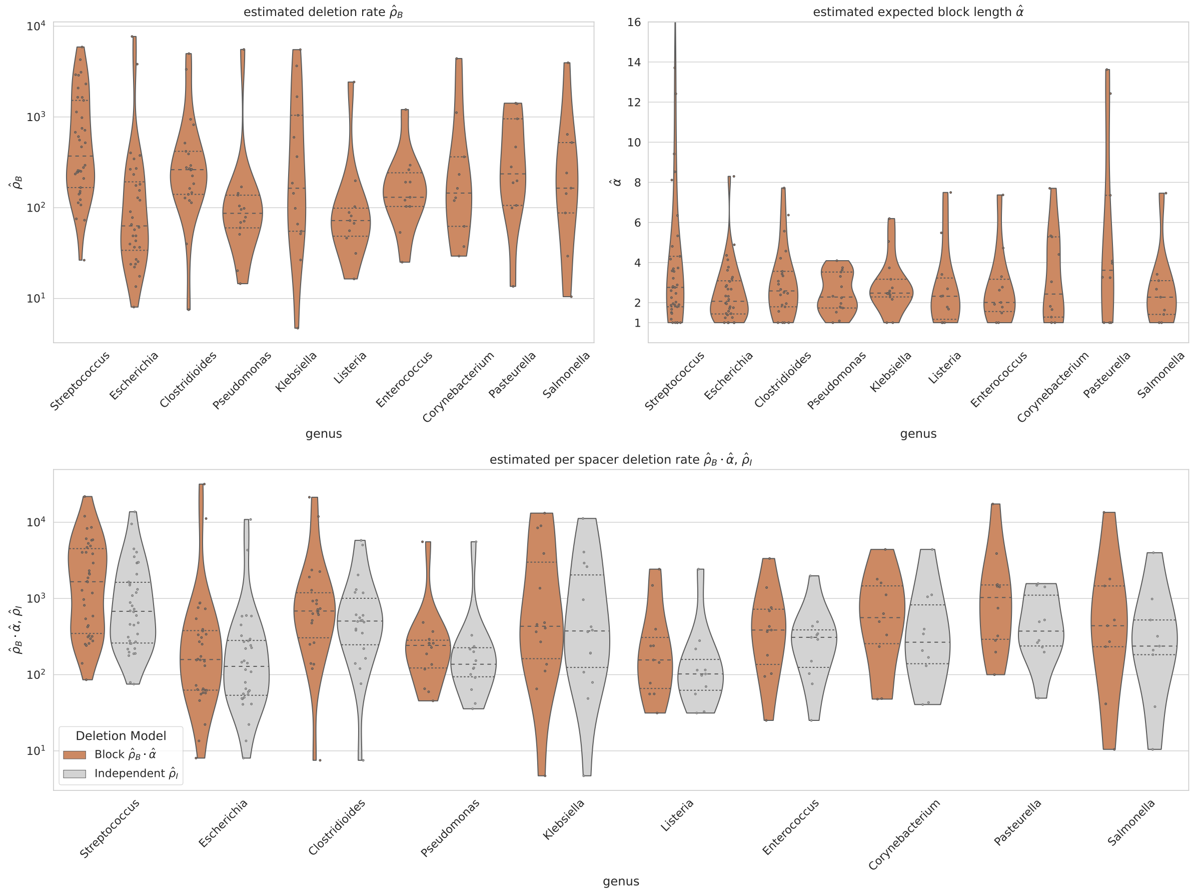


**Supplementary Fig. S16.** We show the estimated parameters for the IDM and BDM of the CRISPRCasdb dataset across prominent genera. We observe no significant deviations between genera, although the differences in the median per spacer deletion rates are larger than across Cas types (see Figure 6 in the main manuscript). This slightly larger deviation might arise due to the differences of mutation rate between the genera. Note, that each groups is composed of samples of the same genus. We only show the estimates for genera with more than 9 available groups with at least one deletion in their reconstruction.

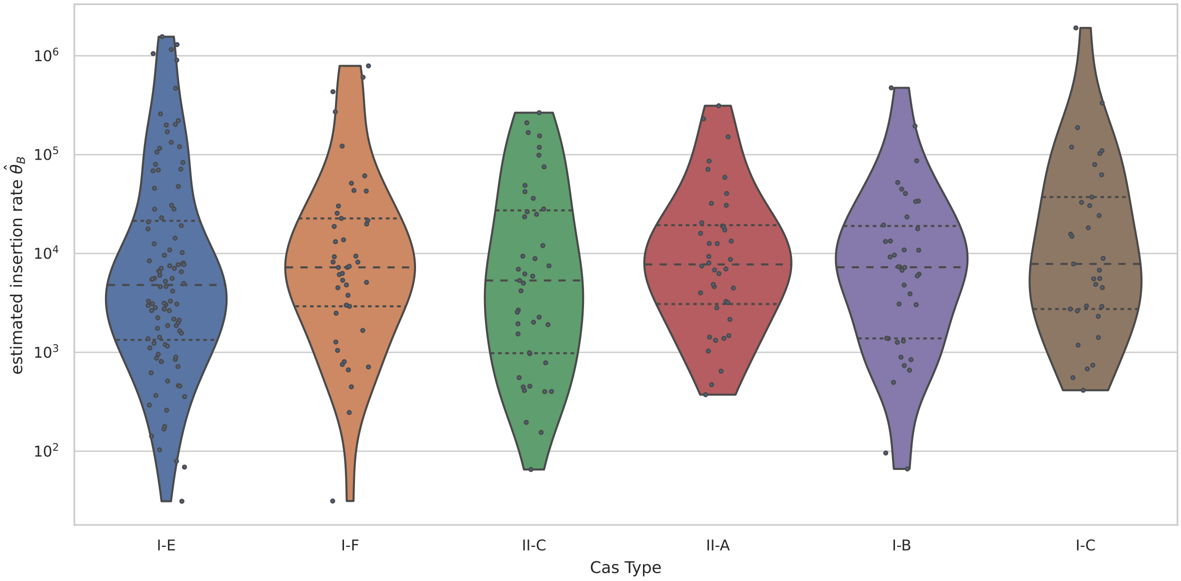


**Supplementary Fig. S17.** We show the estimated insertion rates of the CRISPRCasdb data across prominent Cas types in the dataset. We estimated the insertion rate of each group with the simple estimator $\hat{\theta} = \hat{\rho}_{\mathrm{B}} \cdot \hat{\alpha} \cdot n$, where $n$ is the average array length of the group. Note, that the scale is logarithmic and the variance is quite large. The estimation is unreliable, since this rate is estimated by relying on the reconstructed deletions, which might be sparse or not exist at all. If no deletions were reconstructed, this method is not able to estimate the insertion rate at all. Although, clearly, a group of non-identical arrays acquired spacers at some point. Furthermore, we expect the acquisition process to be much more influenced by interactions with the environment, making average estimates of the insertion rate difficult to interpret.
